# PRC2 facilitates the transition from heterotrophy to photoautotrophy during seedling emergence

**DOI:** 10.1101/2024.10.08.616934

**Authors:** Naseem Samo, María Guadalupe Trejo-Arellano, Lenka Gahurová, Alexander Erban, Alina Ebert, Quentin Rivière, Jiří Kubásek, Fatemeh Aflaki, Helena Hönig Mondeková, Armin Schlereth, Annick Dubois, Mingxi Zhou, Ondřej Novák, Jiří Šantrůček, Daniel Bouyer, Franҫois Roudier, Joachim Kopka, Iva Mozgová

**Affiliations:** Biology Centre CAS, Institute of Plant Molecular Biology, Branišovská 31, České Budějovice, 370 05, Czech Republic; University of South Bohemia in České Budějovice, Department of Molecular Biology and Genetics, Branišovská 31, České Budějovice, 370 05 Czech Republic; Max Planck Institute of Molecular Plant Physiology, Am Mühlenberg 1, Potsdam, 14476, Germany; Laboratory of Growth Regulators, Faculty of Science of Palacký University & Institute of Experimental Botany of the Czech Academy of Sciences, Šlechtitelů 27, CZ-78371, Olomouc, Czech Republic; Laboratoire Reproduction et Développement des Plantes, ENS de Lyon, CNRS, INRAE, Universite Claude Bernard Lyon 1, Lyon, France

**Keywords:** seed-to-seedling, Arabidopsis, heterotrophy, photoautotrophy, Polycomb repressive complex 2

## Abstract

Seed-to-seedling transition represents a key developmental and metabolic switch in plants. Catabolism of seed storage reserves fuels germination and early seedling emergence until photosynthesis is established. The developmental transition is controlled by Polycomb repressive complex 2 (PRC2). However, the coordination of PRC2 activity and its contribution to transcriptional reprogramming during seedling establishment is unknown. By analysing the re-distribution of H3K27me3 and changes in gene transcription in shoot and root tissues of heterotrophic and photoautotrophic seedlings, we reveal two phases of PRC2-mediated gene repression. The first phase is independent of light and photosynthesis and results in irreversible repression of the embryo maturation programme, marked by heterotrophy and biosynthesis of reserve storage molecules. The second phase is associated with the repression of metabolic pathways related to germination and early seedling emergence, and H3K27me3 deposition in this phase is sensitive to photosynthesis inhibition. We show that preventing transcription of the PRC2-repressed glyoxylate cycle gene *ISOCITRATE LYASE* is sufficient to drive the vegetative phase transition in PRC2-depleted plants. This underscores a key role of PRC2 repression in the coordinated metabolic and developmental switches during seedling emergence and emphasizes the close connection between metabolic and developmental identities.

## Introduction

Seedling establishment represents an important developmental and metabolic transition in plants. Seed germination and early seedling growth are fuelled by the catabolism of seed storage reserves before the onset of photosynthesis (Graham, 2008; Tan-Wilson and Wilson, 2012), completing the transition from the heterotrophic to the photoautotrophic growth phase. In *Arabidopsis thaliana* (Arabidopsis), acyl lipids – triacylglycerols (TAGs) are the major seed storage molecules (Li-Beisson et al., 2013). TAGs are broken down by lipases into free fatty acids (FAs) and glycerol (Quettier and Eastmond, 2009). FAs are then converted into acetyl-coenzyme A (Ac-CoA) via β-oxidation (Graham, 2008). Ac-CoA enters the glyoxylate cycle and is converted into 4-carbon compounds by the activities of enzymes including isocitrate lyase (ICL) and malate synthase (MLS) (Graham, 2008). These compounds are then transported to the mitochondria where they can either be processed and transported to the cytosol for gluconeogenesis or used as substrates for respiration.

Seed-stored reserves are used during shoot axis (hypocotyl) elongation following seed germination in the soil (i.e., in darkness), where root growth remains suppressed (skotomorphogenesis). Once the hypocotyl reaches light, its elongation ceases and the cotyledons become photosynthetically active (photomorphogenesis) (Josse and Halliday, 2008; Arsovski et al., 2012). Light signalling and photosynthesis-derived sugars are required to activate the shoot and root apical meristems and to promote cell division and growth of post-embryonic organs (Kircher and Schopfer, 2012; Xiong et al., 2013; Pfeiffer et al., 2016). Seed germination and photomorphogenesis in Arabidopsis involve extensive reprogramming of gene expression (Silva et al., 2016; Narsai et al., 2017; Pan et al., 2023; Tremblay et al., 2024) associated with global and local reorganisation of chromatin (Van Zanten et al., 2012; Bourbousse et al., 2020; Simon and Probst, 2024) and changes in distribution of several histone modifications including H2Bub (Bourbousse et al., 2012), H3K9ac, H3K9me3, H3K27ac, and H3K27me3 (Charron et al., 2009; Pan et al., 2023). Among these, H3K27me3 catalysed by Polycomb Repressive Complex 2 (PRC2) facilitates the seed-to-seedling transition (Holec and Berger, 2012; Mozgova and Hennig, 2015; Hinsch et al., 2021). Deposition of H3K27me3 at the embryo transcription factor genes (*TFs*) *LEAFY COTYLEDON1* (*LEC1*), *LEC1-LIKE, ABSCISIC ACID INSENSITIVE 3* (*ABI3*), *FUSCA3* (*FUS3*), *LEC2* (or collectively “*LAFLs”*) (Gazzarrini and Song, 2024; Aichinger et al., 2009; Bouyer et al., 2011), *AGAMOUS-LIKE 15* (*AGL15*) (Chen et al., 2018) and dormancy regulator *DELAY OF GERMINATION1* (*DOG1*) (Molitor et al., 2014; Chen et al., 2020) leads to their stable repression, facilitating seed germination and seedling establishment. Accordingly, the absence of the PRC2 catalytic subunits CURLY LEAF (CLF) and SWINGER (SWN), or the WD40 subunit FERTILIZATION INDEPENDENT ENDOSPERM (FIE), results in delayed seed germination, failure to develop differentiated tissues, accumulation of TAGs and development of somatic embryos (Chanvivattana et al., 2004; Aichinger et al., 2009; Bouyer et al., 2011; Ikeuchi et al., 2015; Mozgová et al., 2017).

The recruitment of PRC2 to target loci in Arabidopsis is facilitated by distinct families of PRC2-interacting TFs that recognize promoter elements, or Polycomb response elements (PREs), within the target gene promoters (Xiao et al., 2017). Among them, the B3 domain TFs VIVIPAROUS-1/ABSCISIC ACID INSENSITIVE 3-LIKE VAL1/VAL2, which bind the RY element (CATGCA/TGCATG) (Yuan et al., 2021), the telobox-binding TFs TELOMERE REPEAT BINDING PROTEIN1 (TRB1) (Zhou et al., 2018) and ARABIDOPSIS ZINC FINGER 1 (AZF1) (Xiao et al., 2017), and the GA_n_-motif-binding BASIC PENTACYSTEINE (BPC) TFs (Xiao et al., 2017) are responsible for genome-wide PRC2 targeting. *LEC2* PRE-like element is required for the deposition of H3K27me3 and *LEC2* repression in vegetative seedlings (Berger et al., 2011). VAL1 and VAL2 recruit components of the PRC2 to the promoter of *AGL15* (Chen et al., 2018) and to *DOG1* (Chen et al., 2020), facilitating their repression. Beyond this, the dynamics of H3K27me3 deposition and its regulation during seedling emergence is largely unknown. During seed germination and early seedling emergence, H3K27me3 demethylase RELATIVE OF EARLY FLOWERING 6 (REF6) maintains a demethylated state at seedling establishment genes, promoting their expression and seed germination (Pan et al., 2023). Genes encoding PRC2 subunits are transcriptionally activated after seed imbibition (Mozgová et al., 2017; Pan et al., 2023) and changes in H3K27me3 distribution are initiated at 24 - 48 hours (hrs) after seed imbibition (Pan et al., 2023). The nuclear activity of PRC2 is promoted by the kinase TARGET OF RAPAMYCIN (TOR) (Ye et al., 2022), an important integrator of metabolic and nutrient state that balances cell division and growth versus stress response (Ryabova et al., 2019; Wu et al., 2019). However, it remains unknown how the activity of PRC2 and reprogramming of H3K27me3 during seedling emergence alters gene expression to achieve a stable developmental and metabolic transition during seedling establishment.

We show that PRC2 is required to stably repress heterotrophic growth and initiate photoautotrophic (vegetative) growth and development. By profiling the transcriptome and H3K27me3 distribution in wild type (WT) and PRC2-catalytic double mutant seedlings *clf swn* (*cs*) at two metabolically distinct developmental timepoints, we identified two phases of H3K27me3 deposition that underlie a stable transition to vegetative growth. In the first phase, H3K27me3 deposition is independent of light and photosynthesis. It ensures the repression of genes that contribute to a sugar-inducible activation of developmental and metabolic pathways involved in embryo maturation. In the second phase, metabolic processes associated with seed germination/early seedling establishment and early light responses are repressed. The deposition of H3K27me3 at this stage is sensitive to the photosynthesis inhibitor DCMU. By preventing overexpression of the PRC2-target *ICL* in *cs*, we show that PRC2-governed transcriptional repression of metabolic genes during germination is required to promote the developmental transition from seed to seedling.

## Results

### Shoot and root tissues of seedlings before and after the onset of carbon assimilation show different transcriptome and H3K27me3 distribution profiles

To establish the timepoint of transition from heterotrophy to autotrophy, we first estimated the onset of photosynthetic carbon assimilation in emerging seedlings. To do this, we utilized the difference in CO_2_ isotopic composition between atmospheric air during parental plant growth and seed production (δ ^13^C = -8.5‰) and artificial air applied during seed germination (δ^13^C = -40‰) (Šantrůček et al., 2014). Relative decrease of ^13^C showed that photosynthetic CO_2_ assimilation begins after 3 and before 5 days after germination is induced (dag) (i.e., after the transfer of stratified seeds to long-day cultivation) **(Figure 1A**). To understand the PRC2-dependent changes connected to the metabolic transition in source and sink tissues, we analysed gene transcription (**Figure 1B, C** and **Tables S1-S3**) and genome-wide distribution of H3K27me3 (**Figure 1B, D** and **Figure S1A, B** and **Tables S4 - S6**) in the shoots and roots of 3- and 7-dag seedlings. These timepoints represented steady-state heterotrophic and photoautotrophic stages with similar developmental complexity, that allowed manual dissection of shoot (source) and root (sink) tissues. To analyse the effect of heterotrophic growth in the emerging seedlings and identify genes marking heterotrophic growth, we included control plants treated with the photosynthetic inhibitor 3-(3,4-dichlorophenyl)-I,I-dimethylurea (DCMU) that blocks the electron flow from photosystem II to plastoquinone (van Rensen and van Steekelenburg, 1965; Pfister and Arntzen, 1979) (**Figure 1B**). 1% sucrose was added to the DCMU-grown 7-dag seedlings to retain viability.

**Figure 1.**
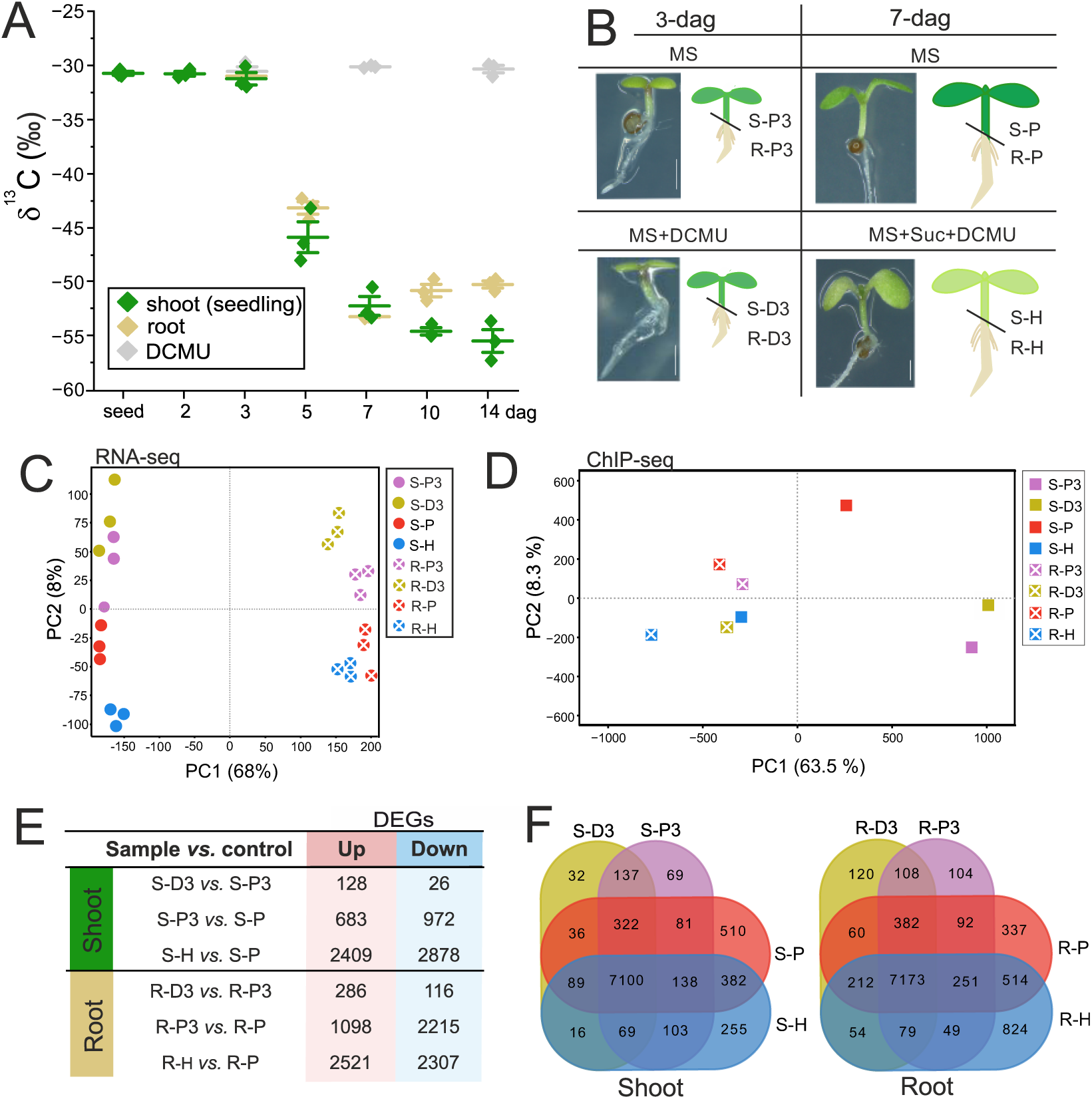
3- and 7-day seedlings display distinct gene transcription and H3K27me3 distribution patterns. **A)** Assimilation of atmospheric CO_2_ is initiated after 3 days after germination initiation (dag). Plants were cultivated in artificial air (δ^13^C = -40‰). Roots and shoots were analysed separately from 3 dag. DCMU: photosynthesis-inhibited control. Each point represents a biological replicate; error bars: ±SD. **B)** Schematic representation of experimental setup. Wild-type seedlings were cultivated on growth medium (MS) supplemented with DCMU (MS+DCMU), or DCMU + 1 % sucrose (MS+Suc+DCMU). Scale bar: 1 mm. Tissue samples: S – shoot; R – root; Cultivation conditions: P – photoautotrophic; H – heterotrophic; D – DCMU; 3 – 3-dag seedling; 7 – 7-dag seedling. **C)** RNA-seq principal component analysis (PCA) plot: RPKM values of all genes. **D)** ChIP-seq PCA plot: genome-wide H3K27me3/H3 values in 200-bp windows, biological duplicates were combined. **E)** Numbers of differentially expressed genes (DEGs) in anaysed sample comparisons. **F)** Overlap of genes enriched for H3K27me3 (H3K27me3 targets) in the shoot and root samples.

Principal component analyses (PCA) of the transcriptomic (**Figure 1C**) and H3K27me3 distribution data (**Figure 1D**, **Figure S1A**) separated shoot and root tissues (PC1) as well as 3- and 7-dag samples (PC2). Heterotrophic growth had a smaller effect on transcription and H3K27me3 distribution in the shoot at 3 dag than at 7 dag (**Figure 1C-D**), indicating lower sensitivity of the 3-dag plants to photosynthesis inhibition. Notably, the PCA of the H3K27me3 distribution (**Figure 1D**, **Figure S1A**) grouped the 7-dag heterotrophic shoot samples with the root samples, indicating a strong effect of the DCMU treatment in 7-dag shoot and shift towards sink-tissue identity. Next, we identified differentially expressed genes (DEGs) between selected pairs of samples (**Figure 1E** and **Tables S2, S3**) and genes targeted by H3K27me3 in each sample (**Figure 1F** and **Tables S5, S6**). High level of H3K27me3 marking (top 25%) over a large proportion of the gene body (≥ 70 %; “T70” targets) was associated with transcriptional repression, while lower H3K27me3 enrichment or smaller proportion of gene body enriched in H3K27me3 were not predictable indicators of transcriptional repression (**Figure S1C - E**). These initial analyses confirmed differences in gene transcription and H3K27me3 distribution between shoot and root tissues in 3- and 7-dag seedlings and underscored the different extent of transcriptional and H3K27me3 remodelling in response to photosynthetic inhibition in the shoot of 3- and 7-dag seedlings.

### Developmental and metabolic genes are repressed in the shoot, while photosynthesis-related genes are repressed in the root of the emerging seedlings

To understand the changes between 3 and 7 dag, we analysed H3K27me3 distribution and gene expression at these two time points in both shoots and roots. To identify genes that require PRC2 for their repression, we first focused on genes that were down-regulated and gained H3K27me3 between 3 and 7-dag. 683 genes were down-regulated between 3 and 7-dag in the shoot (**Figure 1E**, **Table S2**), enriched for plant-type cell wall organisation, lipid catabolism and oxidation-reduction processes (**Table S2**). 1868 PRC2 target genes gained H3K27me3 in the shoot (**Figure 2A** and **Table S7**). 97 of these genes are TF genes (*TFs*) enriched for phyllotactic patterning and negative regulation of flowering, and include the *APETALA 2* (*AP2*) *TFs PLETHORA 3* (*PLT3*), *PLT5/EMK*, *PLT7*, the MADS TF *AGAMOUS-LIKE 15* (*AGL15*), or the flowering repressors *FLOWERING LOCUS C* (*FLC*), *MADS AFFECTING FLOWERING 4* (*MAF4*) and *MAF5*. Other genes that gain H3K27me3 include for instance the auxin transporter *PIN-FORMED 1* (*PIN1*), thylakoid-bound *EARLY LIGHT-INDUCIBLE PROTEIN 1* (*ELIP1*) and *ELIP2*, the glyoxylate cycle enzyme-encoding gene *ICL*, the gluconeogenesis-related enzyme-encoding gene *PHOSPHOENOLPYRUVATE CARBOXYKINASE 1* (*PCK1*) and genes encoding enzymes of reserve lipid catabolism, such as *OIL BODY LIPASE 1* (*ATOBL1*) (**Figure 2B, C** and **Figure S2A** and **Table S7**). This is reflected in decreasing transcription of these genes, including *FLC, ELIP1, PLT5, ICL* and *PCK1* (**Figure 2D**). 261 genes marked by H3K27me3 at 7-dag were downregulated between 3 and 7-dag (**Figure 2E**, **Table S8**). These genes were enriched for biological processes related to epidermis and root development, lipid and hydrogen peroxide catabolism, and oxidative stress response (**Figure 2E**). Notably, the *LAFL* genes were already marked by H3K27me3 and repressed at 3 dag, but further reduction in *ABI3* transcription could still be detected between 3 and 7 dag in the shoot (**Figure 2B - D**). In contrast, 972 genes were upregulated in the shoot between 3 and 7 dag (**Figure 1E**, **Table S2**), of which 375 were marked by H3K27me3 at 3 dag (**Figure S2B**, **Table S8**). These genes were enriched for abiotic and biotic stress responses, hormone signalling including SA and JA, and glucosinolate metabolism (**Figure S2B**). 806 genes lost H3K27me3 between 3 and 7 dag (**Figure 2A** and **Table S7**), 89 of which were *TFs*. These included lateral boundary and shoot organ development *TFs*, such as *NGATHA 2* (*NGA2*) and *NGA3*, TALE homeodomain *TFs BELL1*, *BEL1-LIKE HOMEODOMAIN 2* (*BLH2*), *BLH4*, *KNOTTED1-LIKE HOMEOBOX GENE 4* (*KNAT4*), MYB domain *TFs LATERAL ORGAN FUSION 1* (*LOF1*) and *LOF2*, TEOSINTE BRANCHED 1, CYCLOIDEA and PCF domain (TCP) *TFs* including *TCP2* (**Figure 2B-D** and **Figure S2A** and **Table S7**).

**Figure 2.**
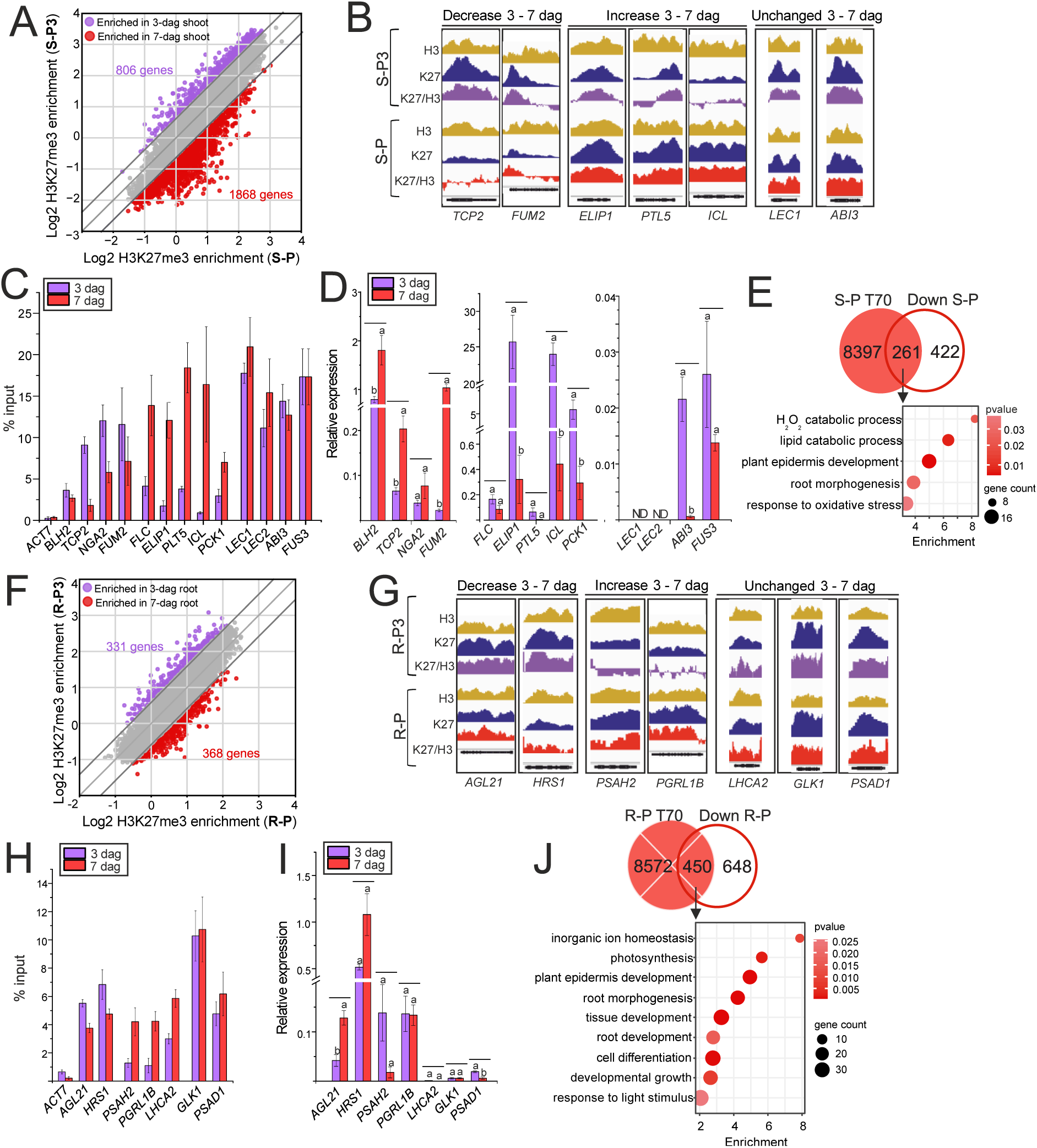
H3K27me3 and transcriptional reprogramming in shoot and root tissues between 3 and 7 dag. A) and. **F)** H3K27me3/H3 enrichment in target genes in 3-dag and 7-dag photoautotrophic shoot (S-P3, S-P; resp.) (A) or root (R-P3, R-P; resp.) (F). Genes enriched for H3K27me3/H3 in at least one sample are displayed. **B) and G)** Genome browser display of selected genes showing the distribution of H3 (ochre), H3K27me3 (blue, “K27”), and H3K27me3/H3 (purple/red “K27/H3”) in 3-dag and 7-dag wild-type shoot (B) or root (G). Y-axis scale of all tracks is identical. **C) and H)** ChIP-qPCR confirmation of ChIP-seq. (C) genes with decreased (*BLH2, TCP2, NGA2, FUM2*), increased (*FLC, ELIP1, PLT5, ICL, PCK1*), and unchanged (*LEC1, LEC2, ABI3, FUS3*) H3K27me3 between 3- and 7-dag shoot. H) genes with decreased (*AGL21* and *HRS1*), increased (*PSAH2* and *PGRL1B*), or unchanged (*LHCA2, GLK1* and *PSAD1*) H3K27me3 between 3- and 7-dag root. *ACT7* serves as negative control with no H3K27me3 enrichment. Bars: mean ±SD; N = 3 technical replicates. **D) and I)** Transcription (RT-qPCR) of genes analysed in C) and H), respectively. Bars: mean ±SD; N = 3 biological replicates. Letters above bars: statistical significance levels at p < 0.01; Student’s *t* test. ND - not detected. **E) and J)** Gene ontology enrichment of 7-dag H3K27me3 target genes (T70) transcriptionally downregulated from 3 to 7-dag in the shoot (E) or root (J). S-P: 7-dag shoot; R-P: 7-dag root. BP categories are shown; GO display cutoff: fold enrichment > 1.5; p(Bonferroni) < 0.05.

Following the approach taken in the shoot, we analysed the differences in H3K27me3 and in the transcriptome between 3- and 7-dag root. 1098 genes were down-regulated between 3 and 7-dag in the root (**Figure 1E** and **Table S3**). These genes were significantly enriched for root morphogenesis, light response and photosynthesis. 368 genes gained H3K27me3 in the root (**Figure 2F-I** and **Figure S2C** and **Table S9**). 450 genes marked by H3K27me3 at 7-dag are downregulated between 3 and 7-dag that are enriched for inorganic ion homeostasis, photosynthesis and light response, as well as root and epidermis development (**Figure 2J** and **Table S10**). Conspicuously, H3K27me3 marked a number of photosynthesis and light response-related genes already at 3 dag (**Figure 2G-I**), indicating an ongoing process that initiated prior to the 3-dag time point. In contrast, 2215 genes activated in the root from 3 to 7 dag were mainly enriched in responses to biotic and abiotic stimuli or stress (**Table S3**). 795 genes marked by H3K27me3 at 3-dag were upregulated between 3 and 7 dag (**Figure S2D**, **Table S10**). Similar to the shoot, these genes were enriched in biotic and abiotic stress responses.

Overall, in both shoots and roots, biotic and abiotic stress response genes were released from H3K27me3-mediated repression and activated between 3 and 7 dag. In the shoot, genes associated with embryo maturation and biosynthesis of embryonic storage compounds were already marked by H3K27me3 and were repressed at 3 dag. Root development and processes associated with seed germination, including catabolism of storage lipids, the glyoxylate cycle and gluconeogenesis, were repressed between 3 and 7 dag. In the root, genes that specifically gained H3K27me3 between 3 and 7 dag and were downregulated during seedling establishment were related to photosynthesis and light responses.

### Heterotrophic growth of 7-dag seedlings reduces the deposition of H3K27me3 and activates photosynthesis-related genes in the root

Heterotrophic growth had a lower impact on H3K27me3 distribution and gene expression in 3-dag seedlings compared to 7-dag seedlings (**Figure 1C - E**). 154 and 402 DEGs were identified in the 3-dag DCMU-grown shoots and roots compared to respective photoautotrophic tissues. In contrast, 5287 and 4828 DEGs, respectively, were identified in 7-dag DCMU-grown shoot and root compared to respective photoautotrophic tissues (**Figure 1E** and **Tables S2, S3**). The genes upregulated in the heterotrophically-grown 7-dag shoot were related to oxidative stress, autophagy and catabolic processes, whereas photosynthesis, anabolic metabolism and SA- and JA-mediated responses were downregulated (**Table S2**). In the heterotrophic root, the upregulated genes were related to stress responses, light response, photosynthesis and chloroplasts, while the down-regulated genes were related to cell division and root development (**Table S3**). Intriguingly, heterotrophically-grown 7-dag seedlings showed a global reduction of H3K27me3 compared to photoautotrophic 7-dag seedlings (**Figure 3A** and **Figure S3A, B**). This was associated with transcriptional increase in the H3K27 demethylases *EARLY FLOWERING 6* (*ELF6*) and the PRC2 catalytic subunit *MEA* in the shoot but no other significant and consistent transcriptional changes of H3K27 demethylases or PRC2 subunits in the shoot or the root (**Figure S3C**). Shoot-specific H3K27me3-target genes were more affected by the H3K27me3 loss than root-specific ones (**Figure 3B** and **Figure S4A - D** and **Table S11**). Genes marked by high levels of H3K27me3 among all the H3K27me3 targets in the shoot and root, and genes that gained H3K27me3 between 3 to 7-dag in the shoot, tended to lose H3K27me3 in heterotrophically-grown seedlings (**Figure 3C, D** and **Table S12**). Genes that lost H3K27me3 in heterotrophically-grown shoots were enriched in development- and transcription-related genes, whereas genes with unchanged H3K27me3 are mostly enriched in metabolism-related Gene Ontology (GO) categories (**Figure S4E** and **Table S12**). Importantly, there was a limited overlap between genes that lost H3K27me3 and genes that were upregulated in heterotrophic shoot compared to photoautotrophic shoot (**Figure S4F**), indicating uncoupled effects of heterotrophic growth on H3K27me3 and gene transcription.

**Figure 3.**
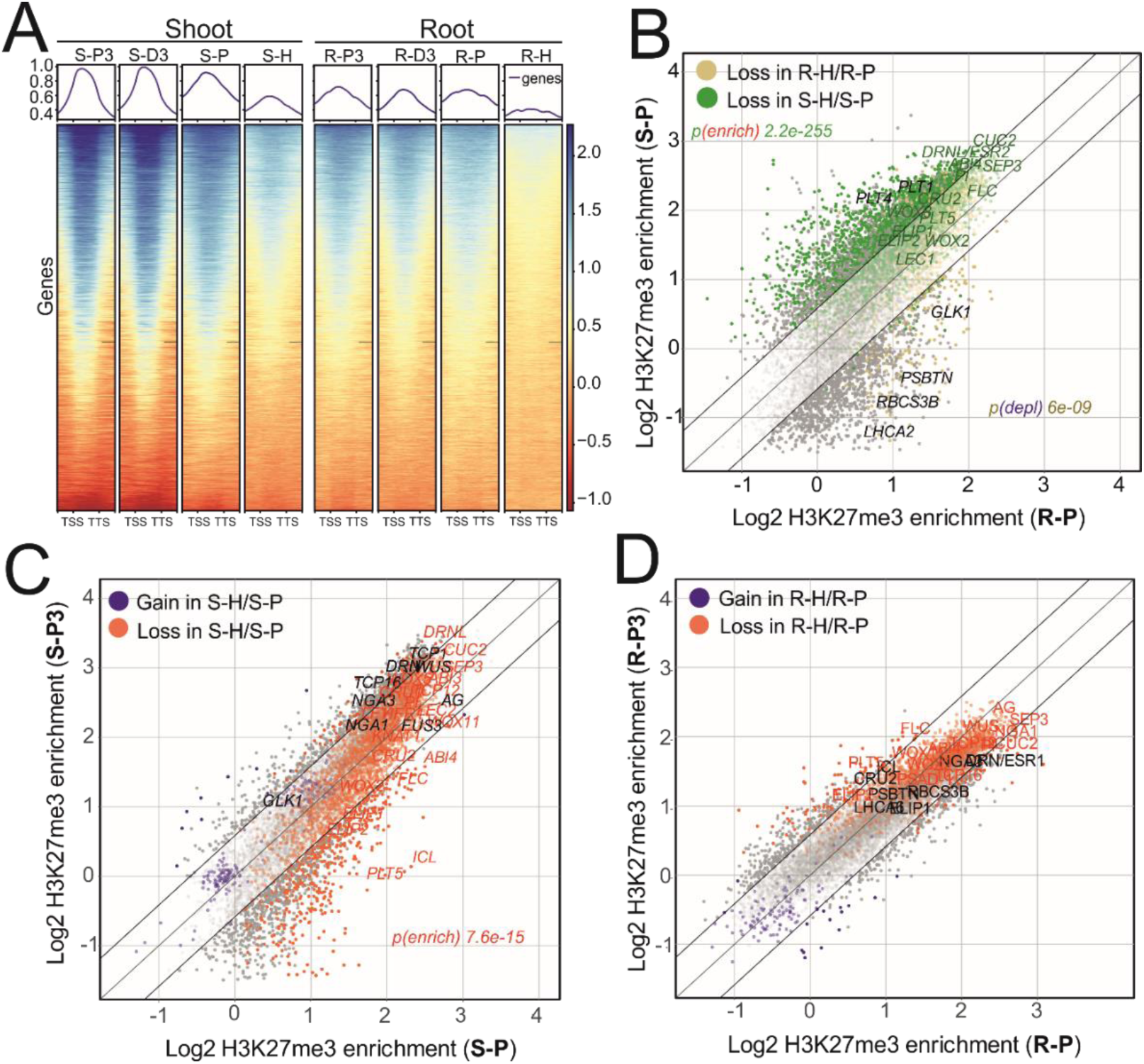
Heterotrophic growth reduces H3K27me3 in 7-dag tissues. **A)** H3K27me3/H3 enrichment in gene bodies (-/+ 0.6 kb) of H3K27me3 target genes in wild-type shoot or root. Plotted are H3K27me3 target genes (T70) identified in at least one of the shoot samples (“Shoot” panel) or root samples (“Root” panel). Sample labels correspond to Figure 1B. **B)** H3K27me3/H3 enrichment in H3K27me3 target genes (T70) in 7-dag shoot (S-P) and root (R-P). Each dot represents a gene; green and ochre dots represent genes that lose H3K27me3 in 7-dag heterotrophic shoot (S-H/S-P) and root (R-H/R-P), resp. p-values: significance of overlap (enrichment or depletion) between shoot (green) or root (ochre) genes and genes losing H3K27me3 in respective heterotrophic samples. **C)** H3K27me3/H3 enrichment in H3K27me3 target genes (T70) in 3-dag (S-P3) and 7-dag (S-P) photoautotrophic shoot. Each dot represents a gene; blue or red dots represent genes that gain or lose H3K27me3, respectively, in 7-dag heterotrophic compared to photoautotrophic shoot (S-H/S-P). p-values: significance of overlap between genes that lose H3K27me3 in heterotrophic shoot (S-H/S-P) and genes that gain H3K27me3 between 3 to 7 dag. **D)** H3K27me3/H3 enrichment in H3K27me3 target genes (T70) in 3-dag (R-P3) and 7-dag (R-P) photoautotrophic root. Each dot represents a gene; blue or red dots represent genes that gain or lose H3K27me3, respectively, in 7-dag heterotrophic compared to photoautotrophic root (R-H/R-P).

### PRC2 represses exogenous sugar-induced accumulation of storage lipids and promotes photoautotrophic growth

Embryonic depletion of FIE (Bouyer et al., 2011) or CLF and SWN (**Figure S5A - C**) does not impede embryo or seed development, but it significantly affects postembryonic development (**Figure 4A**). Germination of *cs* seeds was delayed by 2 – 3 days regardless of growth conditions (**Figure S5D**) and seedlings developed cotyledon-like pale green shoot structures that accumulated TAGs (**Figure 4A** – “*cs*-M” and **Figure S5E, F**) (Chanvivattana et al., 2004; Aichinger et al., 2009; Bouyer et al., 2011). Importantly, the described developmental phenotypes and the ectopic accumulation of storage lipids was conditioned by continuous presence of sucrose in the cultivation medium since germination, or its supply before 3 dag (**Figure S5G, H**). In contrast, in the absence of exogenous sucrose or its supply after 3 dag, green vegetative *cs* plantlets developed that did not accumulate TAGs (**Figure 4A** - “*cs*-P” and **Figure S5E - H**). Unlike WT plants, sucrose-grown *cs* plants failed to assimilate atmospheric CO_2_ (**Figure 4B**) but accumulated biomass (**Figure S5I**), indicating heterotrophic growth. In contrast, *cs* plants grown without external sucrose assimilated atmospheric CO_2_ (**Figure 4B**) and accumulated biomass (**Figure S5I**), indicating photoautotrophic growth. Notably, the assimilation and growth rate in photoautotrophic *cs* plants was significantly lower than in WT plants and they exhibited embryonic flower-like phenotypes and homeotic defects (**Figure 4A** – “*cs*-P”). These results led us to conclude that PRC2 activity in the initial phases of seedling emergence is essential to repress pathways that direct sucrose towards accumulation of storage lipids and to stimulate photoautotrophic growth.

**Figure 4.**
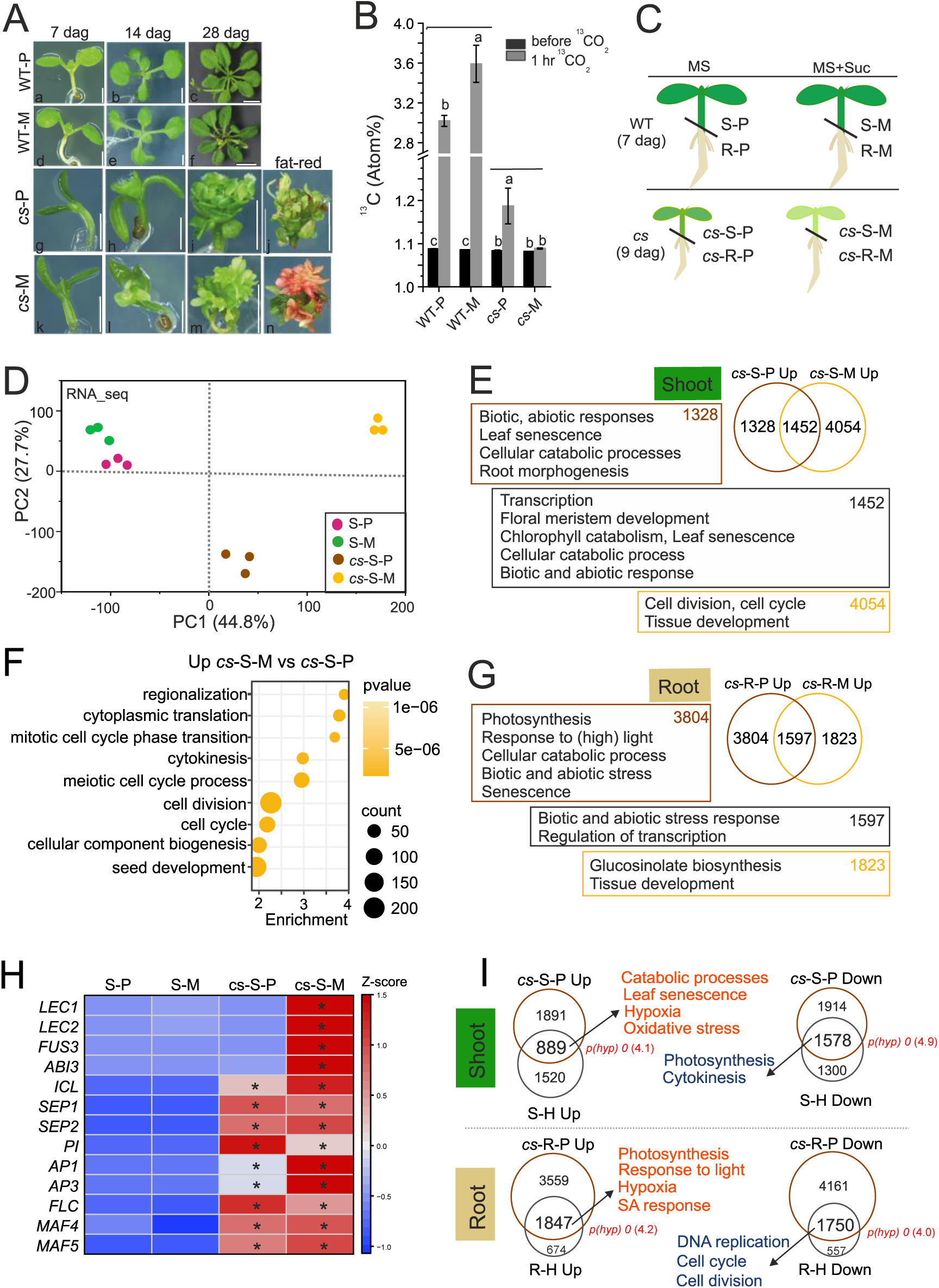
PRC2 is required for photoautotrophic growth. **A)** Wild type (WT; a-f) and *clf swn* (*cs*; g-n) plants cultivated in photoautotrophic (P) and mixotrophic (M) conditions for 7, 14 and 28 days. Fat red staining (j, n) is used to detect embryonic lipids. Scale bar = 1 mm (a, d, g, h, k, l), 2 mm (i, j, m, n), 5 mm (b, e), 1cm (c,f). **B)** ^13^CO2 content in the shoot before and after 1-hr cultivation in ^13^CO_2_-enriched air. WT and *cs* shoots of 10-dag seedlings cultivated in photoautotrophic (P) and mixotrophic (M) growth conditions are shown. Approximately 1.1 atom% of ^13^C in all WT and *cs* represents natural abundance of ^13^CO_2_ in atmospheric air. Bars: mean ±SD; N = 3 biological replicates. Letters above bars: p < 0.05; two-way ANOVA with Bonferroni post hoc test. **C)** Experimental setup of transcriptome analyses of seedlings 5 – 6 days after germination. 9-dag *cs* and 7-dag WT is used to account for the delay in *cs* germination. S – shoot; R – root; P – photoautotrophic; M – mixotrophic; MS – ½ Murashige and Skoog medium; suc - sucrose. **D)** RNA-seq principal component analysis (PCA) plot: RPKM values of all genes. **E)** Schematic representation of genes and enriched GO biological processes upregulated in *cs* photoautotrophic (*cs*-S-P) and mixotrophic (*cs*-S-M) shoots compared to respective WT shoots (S-P and S-M). GO summary cutoff: p(Bonferroni) < 0.05. **F)** Biological processes upregulated in mixotrophic (*cs*-S-M) compared to photoautotrophic (*cs*-S-P) *cs* shoot. GO summary cutoff: p(Bonferroni) < 1e-06. **G)** Schematic representation of genes and enriched GO biological processes upregulated in *cs* photoautotrophic (*cs*-R-P) and mixotrophic (*cs*-R-M) roots compared to respective WT roots (R-P and R-M). GO display cutoff: p(Bonferroni) < 0.05. **H)** Expression of embryo and flower development genes in WT and *cs*. Z-scored RPKM; asterisks (*): significantly different transcription related to corresponding WT (FDR < 0.05, DESeq2). **I)** Photoautotrophic *cs* seedlings resemble heterotrophic WT seedlings. p(hyp): p-value - hypergeometric test of enrichment; ratio of observed/expected indicated in brackets. Summary of GO biological processes enriched among genes commonly dysregulated in photoautotrophic *cs* and DCMU-treated WT tissues. GO summary cutoff: fold enrichment > 3; p(Bonferroni) < 0.05. Full GO graphs are shown in Figure S7.

To identify transcriptional patterns associated with phenotypic differences in *cs* (**Figure 4A**), we analysed the transcriptome of photoautotrophic and mixotrophic WT and *cs* shoots and roots 5 - 6 days after the seeds have germinated (i.e., after the switch to photoautotrophy - **Figure 1A**). To account for the delayed germination in *cs*, 7-dag WT and 9-dag *cs* plants were analysed (**Figure 4C** and **Figure S6A** and **Table S13**). Consistent with the observed differences in developmental phenotypes (**Figure 4A**), sucrose enhanced the separation of WT and *cs* shoot samples, and the transcriptome of WT was less affected by the presence of sucrose than that of *cs* (**Figure 4D**). This was reflected in 739 sucrose-induced DEGs in WT but 10896 in *cs* shoot (**Figure S6B** and **Table S13**). Similar effect was observed in the root tissues (**Figure S6A** and **Table S14**), corroborating massive sucrose-induced transcriptional reprogramming in *cs*.

To understand the exogenous sugar-dependent or independent effects of *cs* on transcription, we compared DEGs in *cs* compared to WT in mixotrophic or photoautotrophic shoots (**Figure 4E, F** and **Figure S6C, E** and **Tables S13, S15**). In *cs* shoot (**Figure 4E**), 1452 genes related to transcription, floral meristem, catabolic processes, biotic and abiotic responses and senescence were upregulated in both growth conditions. Similar processes were also enriched among the 1328 genes that were specifically upregulated in the photoautotrophic shoot. 4054 genes that were only upregulated in the mixotrophic *cs* shoot were related to the cell cycle and tissue development. Conversely, photosynthesis and light response were downregulated in the *cs* shoot under both growth conditions (**Figure S6C**). Genes related to cell division and seed development were upregulated in the mixotrophic compared to photoautotrophic *cs* shoot (**Figure 4F**), while biotic and abiotic stress responses and light responses were upregulated in photoautotrophic *cs* shoot (**Figure S6E**). Importantly, upregulation of the *LAFLs* - *LEC1*, *LEC2* and *FUS3* - was limited to mixotrophic *cs* (**Figure 4H**, **Figure S6F**), as was the accumulation of abscisic acid (**Figure S6G**). In contrast, metabolic genes involved in early seedling establishment, including *ICL*, were upregulated regardless of growth conditions. Similarly, other known PRC2-target genes, including the floral identity MADS-box TF genes *SEPALLATA 1* (*SEP1*), *SEP2*, *PISTILLATA* (*PI*), *APETALA 1* (*AP1*), *AP2* or the floral repressor *FLC*, *MAF4* or *MAF5*, were upregulated independently of sucrose, indicating an uncoupled effect of sucrose on the regulatory networks controlling different developmental pathways (**Figure 4H**). In the *cs* root, genes connected to biotic and abiotic stress were upregulated in both conditions, while photosynthesis and light response genes were upregulated mainly in the photoautotrophic root (**Figure 4G**). At the same time, root morphogenesis and growth-related genes were downregulated, especially in photoautotrophic *cs* roots (**Figure S6D**). Notably, the transcriptomic changes in photoautotrophic *cs* resembled heterotrophically-grown WT plants (**Figure 4I** and **Figure S7A - E** and **Table S16**), marked by elevated senescence- and stress-related gene expression in the shoot, induction of photosynthesis and light-response genes in the root, and by general suppression of cell division- and growth-related genes. Collectively, PRC2 represses sucrose-induced upregulation of genes related to seed development, including *LAFL*, ectopic accumulation of storage lipids, and heterotrophic growth. In the absence of exogenous sucrose, PRC2 is not essential for the stable transition to photoautotrophy, but promotes photosynthetic carbon assimilation and vegetative growth.

### Shoot of PRC2-depleted seedlings displays transcriptional and metabolic signatures of developing seeds and early emerging seedlings

Transcriptional changes in *cs* indicated activation of metabolic programs associated with seed maturation, germination and early seedling establishment. Therefore, we compared genes upregulated in *cs* shoot with previously published clusters of genes peaking at different stages of Arabidopsis seed-to-seedling transition (Silva et al., 2016) (**Figure 5A** and **Figure S8A - C**). The photoautotrophic *cs* shoot resembled dry seeds and early greening seedlings, whereas mixotrophic *cs* shoot showed highest similarity to dry and early germinating seeds (**Figure 5A**). Next, we analysed the transcription of key embryo maturation and seedling establishment metabolic genes in WT at the early greening stage (2 dag) and after photoautotrophic transition (7 dag), and compared this to 9-dag *cs*, using photoautotrophic and mixotrophic seedling shoots (**Figure 5B**). 9-dag *cs* resembled 2-dag WT, particularly in mixotrophy that promoted transcription of β-oxidation, glyoxylate metabolism and TCA cycle genes. Importantly, the *LAFL* TFs were not activated by exogenous sucrose even in 2-dag WT, indicating that they were stably repressed at this stage. In *cs*, transcription of *LAFLs* was comparable to 2-dag WT levels in photoautotrophic state but transcripts strongly accumulated in mixotrophic sample, i.e. upon supplementation by external sucrose (**Figure 5B**).

**Figure 5.**
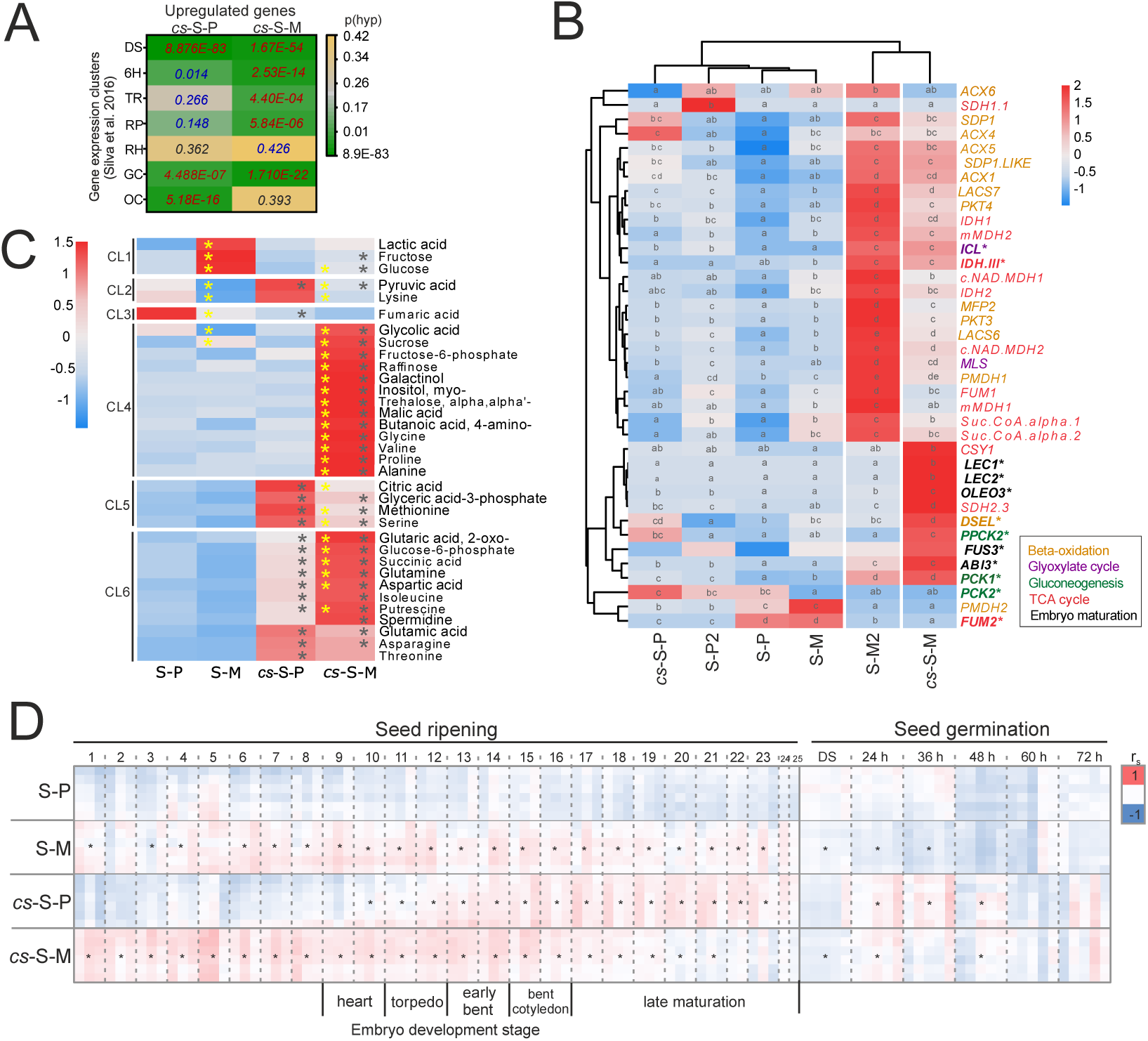
The transcriptome and primary metabolome of PRC2-depleted *clf swn* (*cs*) seedlings resembles embryo maturation and seed germination developmental stages. **A)** Comparison of differentially expressed genes in 9-dag photoautotrophic (*cs*-S-P) and mixotrophic (*cs*-S-M) *cs* seedling shoots to genes transcribed in dry seed and seedling establishment stages(Silva et al., 2016). P-values of hypergeometric tests for gene set overlaps: enrichment (red font) and impoverishment (blue font). DS - dry seed, 6H - six hours of imbibition, TR - testa rupture, RP - radicle protrusion, RH - root hair emergence stage, GC – greening cotyledon stage, OC - open cotyledons stage (cf. full dataset comparison in Figure S8B, C). **B)** RT-qPCR-based determinations of transcript abundances of genes involved in embryo maturation and metabolic pathways marking seedling emergence in photoautotrophic (S-P2 and S-P) and mixotrophic (S-M2 and S-M) WT shoot at 2- and 7-dag and photoautotrophic (*cs*-S-P) and mixotrophic (*cs*-S-M) *cs* shoot at 9-dag. Mutant sampling was delayed to adjust mutant development to 7-dag WT. The heatmap represents z-score normalized relative expression mean of three biological replicates. Letters (a – e): statistical significance at p < 0.05, based on a two-way ANOVA. **C)** Heatmap of selected relative metabolite abundances in WT photoautotrophic (S-P) and mixotrophic (S-M) shoots of WT seedlings compared to respective *cs* shoot samples *cs*-S-P and *cs*-S-M. Z-score normalized means are represented; N = 5 biological replicates. Asterisks (*): statistical significance at p < 0.05 (adjusted) based on ANOVA and Tukey’s HSD test. Black asterisks indicate significant difference among genotypes (*cs*-S-P vs. S-P and *cs*-S-M vs. S-M), yellow asterisks indicate significant differences induced by mixotrophy (S-M vs. S-P and *cs*-S-M vs. *cs*-S-P). Clusters (CL) 1-6 correspond to Figure S8F, i.e., the complete heatmap of all 85 metabolites. **D)** Spearman correlation between the primary metabolomes of photoautotrophic (P) and mixotrophic (M) WT and *cs* shoot samples and samples representing 25 stages of seed maturation, dry seeds (DS) and 5 stages of seed germination sampled at 24, 36, 48, 60 and 72 hours after imbibition(Ginsawaeng et al., 2021). Metabolite data were maximum-scaled per metabolite and dataset to a 0 – 100 numerical range. Negative correlation (-1 minimum correlation coefficient, blue), non-correlated (0, white), and positive correlation (+1 maximum correlation coefficient, red). Asterisks (*): significant differences between replicates of correlation coefficients (Student’s t-test, p > 0.05, 2-tailed, heteroscedastic) tested against the respective WT photoautotrophic (S-P) replicates.

To understand whether these changes are reflected in the metabolome, we analyzed primary metabolites in 7-dag WT and 9-dag *cs* shoots under photoautotrophic and mixotrophic conditions (**Figure 5C** and **Figure S8D-F** and **Table S17**). PCA and contribution plot of the primary metabolome indicated that the differences in genotypes and supplementation of sucrose were the main sources of variation (**Figure S8D, E**). Amino acids and organic acids contributed highly to the differences between genotypes. Sucrose, fructose and glucose among other metabolites distinguished photoautotrophy from mixotrophy (**Figure S8E**). The 85 identified primary metabolites were grouped into 6 clusters (CL1-CL6, **Figure S8F**), highlighting the differences between genotypes and the differential effects of exogenous sucrose on WT or *cs*. In particular, metabolization of sucrose provided to mixotrophic shoot differed between WT and *cs* shoots. WT accumulated fructose (Fru) and glucose (Glc) (CL1) indicating uptake and a limited metabolization of sucrose, e.g., by invertase-catalyzed cleavage. In contrast, *cs* phosphorylated Glc and Fru and metabolized hexose-phosphates further (CL4, CL5, and CL6). Glc6P and Fru6P accumulated in *cs* together with glyceric acid-3P and intermediates of the TCA cycle, including malic acid, citric acid, 2-oxoglutaric acid and succinic acid (**Figure S8F**). Catabolism of supplied sucrose extended up to and beyond the TCA and was associated with anabolic accumulation of most proteinogenic amino acids in photoautotrophic and mixotrophic *cs* (CL5 and CL6). In contrast to WT or photoautotrophic *cs*, mixotrophic *cs* accumulated raffinose, galactinol and myo-inositol, metabolites of the raffinose family oligosaccharide (RFO) biosynthesis pathway, as well as salicylic acid, proline, 4-aminobutyric acid (GABA), and trehalose (CL4). Similarly, putrescine and spermidine accumulated most in mixotrophic *cs* (CL6). The presence of these marker metabolites indicated pronounced physicochemical stress (Dempsey et al., 2011; Zandalinas et al., 2022). Mixotrophic *cs* accumulated TAGs (**Figure 4A** and **Figure S5E - H**). This process is expected to consume acetyl-CoA for fatty acid biosynthesis and to redirect acetyl-CoA from metabolization by the TCA cycle. Indeed, citric acid (CL5) that is synthesized by citrate synthase from acetyl-CoA and oxaloacetate at the entry point of the TCA cycle was the only intermediate of the TCA cycle that accumulated less in mixotrophic *cs* than in photoautotrophic *cs*.

Next, we compared the metabolome of the *cs* shoot with different stages of seed development, maturation and germination. We profiled the metabolome of 25 WT seed developmental stages, corresponding to embryo morphogenesis stages up to the late embryo maturation (**Figure S9A** and **Table S18**). In addition, we analyzed previously published data of primary metabolites from developmental series of germinating seeds (Ginsawaeng et al., 2021). 53 metabolites were robustly identified in all samples and overlapped with the current study (**Table S19**). Weighted correlation network analysis (WGCNA) indicated that relative amounts of amino acids and organic acids are depleted in dry seeds but accumulate during germination (**Figure S9B - E** and **Table S20**), which was consistent with earlier studies (Fait et al., 2006; Silva et al., 2017). Next, we compared the abundance patterns of samples from the current study to the chosen reference profiles by nonparametric correlation across the 53 commonly detected metabolites (**Figure 5D** and **Table S21**). The primary metabolome of the photoautotrophic WT shoot was largely uncorrelated or correlated negatively with the seed maturation or germination developmental samples. In contrast, the samples from photoautotrophic *cs* shoot correlated positively with the late maturation stages of embryogenesis and seedlings at 24 - 48 hrs after germination. Exogenous sucrose in WT induced metabolic changes that enhanced correlations to the early to intermediate stages of seed development. The positive correlation with early seed developmental stages was further enhanced in *cs* samples (**Figure 5D**).

In summary, *cs* seedlings exhibited transcriptional and metabolic characteristics of maturing embryos, germinating seeds, and early seedling establishment stages. In particular, transcripts of genes involved in the degradation of storage lipids, i.e. fatty acid β-oxidation, the glyoxylate cycle, gluconeogenesis and the TCA cycle accumulated. These transcriptional similarities are associated with the metabolic signatures of late embryogenesis and germinating seeds that are retained in *cs* seedlings. Application of exogenous sucrose promotes metabolic patterns of early developing embryos in *cs* seedlings beyond the WT and induces a complex metabolic stress response.

### Two phases of PRC2 repression are required for the establishment of photoautotrophic seedling

Based on the preceding analyses, we hypothesised that before 3 dag, PRC2 represses pathways that direct sucrose towards TAG biosynthesis and promote embryo development. To identify the underlying genes, we compared genes marked by H3K27me3 at 3-dag with genes upregulated in mixotrophic, but not in photoautotrophic, *cs* shoot (**Figure 6A**). We identified 564 genes enriched for lipid storage and associated GO categories. Among these were 82 *TFs*, including *LEC1*, *LEC2* and *FUS3* (**Figure 6A** and **Table S22**). Next, we asked which pathways require PRC2 for their repression in order for the vegetative (photoautotrophic) state to be established. 805 genes marked by H3K27me3 in 3- and 7-dag

**Figure 6.**
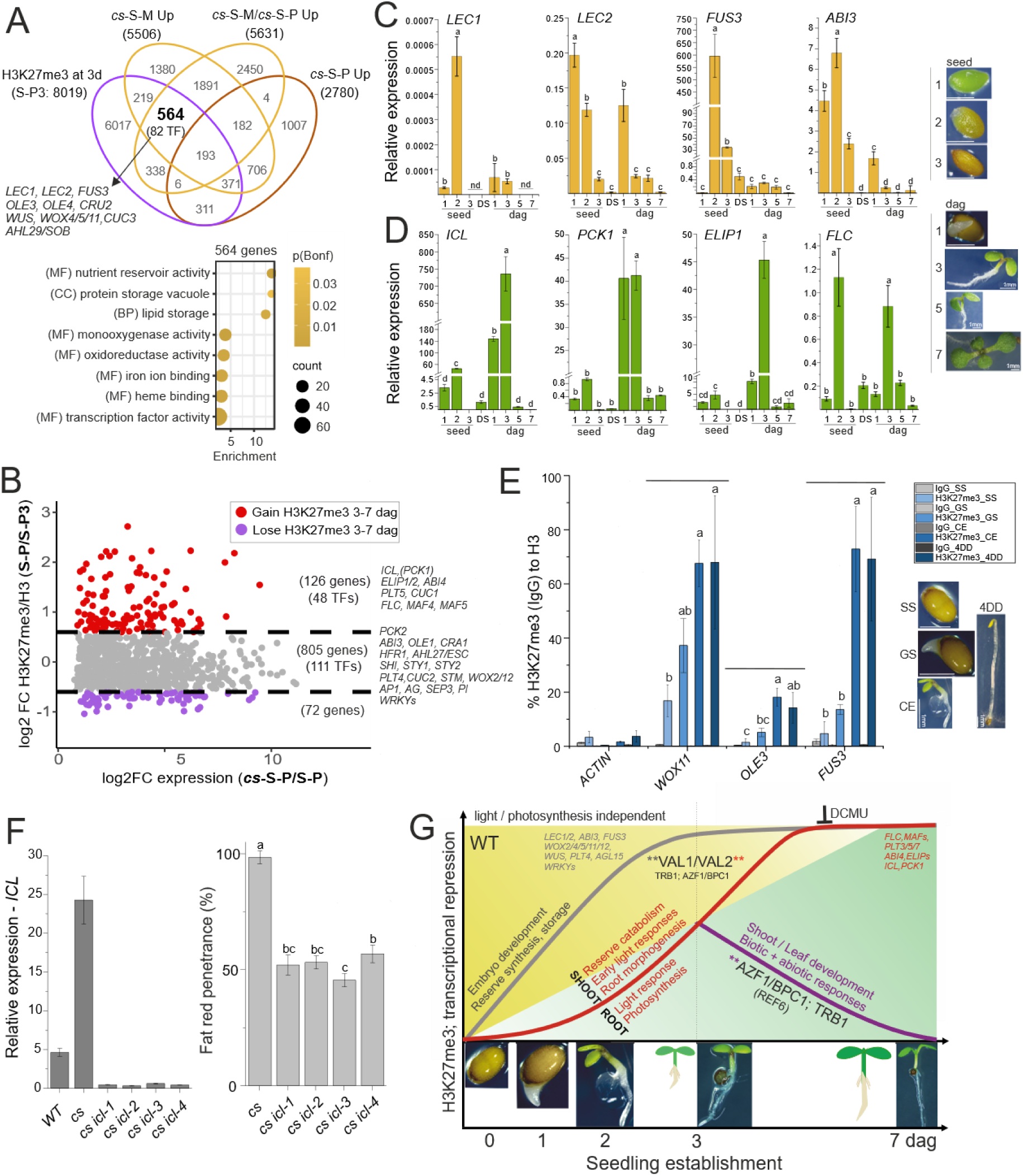
PRC2 coordinates developmental and metabolic reprogramming in several phases of seedling establishment. **A)** Identification of H3K27me3-target genes that contribute to reversal to lipid accumulating (embryo maturation) phase in mixotrophically-grown *clf swn* seedlings. Comparison of gene sets marked by H3K27me3 at 3-dag (S-P3), genes upregulated in mixotrophic *cs* shoot compared to respective WT (*cs*-S-M Up), genes upregulated in photoautotrophic *cs* shoot compared to respective WT (*cs*-S-P Up) and gene upregulated in mixotrophic compared to photoautotrophic *cs* shoot (*cs*-S-M/*cs*-S-P Up). Selected key regulators of embryo development are highlighted among 564 protein-coding genes. GO analysis of 564 genes that contribute to the metabolic/developmental reversal. GO display cutoff: fold enrichment > 1.5; p(Bonferroni) < 0.05. **B)** Identification of genes marked by H3K27me3 at 3 and/or 7-dag that contribute to the *cs* shoot phenotype during photoautotrophic growth. Y-axis: fold-change (FC) H3K27me3/H3 enrichment in 7-dag (S-P) compared to 3-dag (S-P3) WT shoot; X-axis: FC expression of DEGs upregulated in *cs* (*cs*-S-P) compared to WT (S-P) photoautotrophic shoot. Expression log2 FC cutoff is 0.6. Each dot represents a gene: red dots: 126 genes (48 transcription factor genes -TFs) that gain H3K27me3 between 3- and 7-dag WT shoot and are upregulated in *cs*; purple dots: 72 genes that lose H3K27me3 between 3- and 7-dag WT shoot and are upregulated in *cs*; grey dots: 805 genes (111 TFs) with unchanged levels of H3K27me3 between 3- and 7-dag WT shoot. H3K27me3/H3 log2 FC cut-off is 0.6. Selected key developmental or metabolic regulators are highlighted. **C)** Expression of genes marked with H3K27me3 by 3 dag. **D)** Expression of genes gaining H3K27me3 between 3 and 7 dag. **C)** and **D)** qRT-PCR analysis of selected gene transcription in 3 seed development stages (“seed”: 1 – mature green, 2 – mature yellowing, 3 – mature desiccating), dry seeds (DS) and 4 stages of seedling germination (dag: 1, 3, 5 and 7). Representative seeds/seedlings are shown on the right. Bars: mean ± SD; N = 3 biological replicates. Letters above bars: statistical significance at p < 0.05; one-way ANOVA with Bonferroni post hoc test. ND - not detected. **E)** H3K27me3 deposition during seed germination is independent of active photosynthesis. ChIP-qPCR of H3K27me3 enrichment at four stages of seedling establishment before the onset of photosynthesis: stratified seed (SS), germinated seed (GS), cotyledon emergence (CE) and 4-dag dark-grown seedling (4DD). Bars: mean ± SD; N = 3 biological replicates. Letters above bars: statistical significance at p < 0.05; one-way ANOVA with Bonferroni post hoc test. **F)** CRISPR-Cas9 mutagenesis of *ICL* in *clf swn* genetic background (*cs icl*) limits the embryonic reversal in *cs*. RT-qPCR of *ICL* expression (left). Bars: mean ± SD; N = 3 technical replicates. Penetrance of positive fat-red staining phenotype (right) in *cs (icl)* seedlings grown in the presence of 1% sucrose. Bars: mean ± SD; N = 3 biological replicates (20 - 30 *cs icl* seedlings/replicate). **G)** Summary of PRC2 contribution to gene repression at different phases of seedling establishment. Backgrounds represent seed (heterotrophic - yellow) to seedling (photoautotrophic – green) transition. Grey and red lines/font represent increase in H3K27me3 (transcriptional repression) before 3-dag and between 3 - 7 dag, respectively. Purple line/font represent decrease in H3K27me3 (transcriptional activation) between 3 – 7 dag. Representative developmental seed and seedling stages are shown in the bottom.

WT shoot are upregulated in photoautotrophic *cs* compared to WT shoot (**Figure 6B** and **Table S23**). These genes were involved in seed, root or flower development, auxin biosynthesis, oxidative stress response and biotic defence responses (**Table S23**). 111 genes encode TFs, including key transcriptional regulators of these processes (**Figure 6B**). 126 (48 TFs) genes upregulated in photoautotrophic *cs* shoot gained H3K27me3 between 3 and 7-dag in WT (**Figure 6B** and **Table S23**). This group of genes included genes related to the glyoxylate cycle (*ICL*) and gluconeogenesis (*PCK1*), seed development (*ABI4*, *LEAs*), early light response (*ELIP1*, *ELIP2*), root (*PLT5*) and shoot (*CUC1*) development and the flowering repressors *FLC*, *MAF4* and *MAF5*. Hence, PRC2 transcriptionally repressed distinct groups of genes during the seed-to-seedling transition – those repressed by 3 dag, potentially reactivated by exogenous sucrose, and those repressed between 3-7 dag. Accordingly, the transcription of seed development genes peaked during embryo maturation and at 1 dag, but was repressed at 3-dag (**Figure 6C**). In contrast, transcription of seed germination metabolic genes peaked at 3-dag and was repressed by 7-dag (**Figure 6D**). Activation of the *LAFL* transcriptional network in mixotrophic *cs* resembled *LEC1*-overexpressing plants (Mu et al., 2008), but was independent of activation of fatty acid catabolism and glyoxylate cycle-associated genes, limited to *cs* (**Figure S10A** and **Table S24**). These observations suggested that the pathways of seed maturation and seed germination are independently regulated, but all are simultaneously activated in mixotrophic *cs*. We next asked whether repression of seed development genes before 3-dag requires post-germination PRC2 activity or whether H3K27me3 is established in seeds. We found that the amount of H3K27me3 increased at these loci following seed imbibition until 2–3-dag, and this increase was also detected in dark-grown etiolated seedlings (**Figure 6E**). To determine if the identified sets of genes repressed by 3 and 7-dag may differ in the mode of PRC2 recruitment, we analysed the enrichment of VAL1/VAL2-, TRB1- and AZF1/BPC1-target genes and known PRE motifs among the identified subsets of H3K27me3 targets (**Figure S10B - D**). We found that genes marked by H3K27me3 at 3-dag but not at 7-dag (378 genes) were enriched in TRB1 and AZF1/BPC1 targets (**Figure S10C**), and their promoters were enriched for the REF6-binding motif (CTCTGTT) (**Figure S10D**). Genes upregulated in photoautotrophic *cs* shoot with comparable H3K27me3 marking at 3 and 7-dag (805 genes) were enriched in all, VAL1/VAL2-, TRB1- and AZF1/BPC1-targets and the VAL1/VAL2-bound RY-motif (CATGCA). In contrast, H3K27me3 targets induced by sucrose in *cs* (564 genes) and genes that gained H3K27me3 between 3 and 7-dag (126 genes) were significantly enriched only in VAL1/VAL2 targets (**Figure S10C**) and the RY-motif in the case of the genes gaining H3K27me3 between 3 and 7-dag (126 genes) (**Figure S10D**).

Having identified metabolic genes that gain H3K27me3 during seedling establishment, we asked whether PRC2-mediated repression of metabolic pathways is needed to establish photoautotrophic growth. We used CRISPR/Cas9 in *cs* to knock out the key glyoxylate cycle gene *ICL*, a single-copy gene targeted by H3K27me3 and transcriptionally repressed between 3 – 7-dag, that is upregulated in *cs*. We hypothesised that ectopic activation of *ICL* may prevent greening and seedling establishment, in which case *cs icl* triple mutants should green regardless of growth conditions. Indeed, several independent alleles of *cs icl* showed a significantly reduced penetrance of the TAG-accumulating phenotype and a higher frequency of transition to the vegetative phase (**Figure 6F** and **Figure S10E- I**). These data demonstrated that PRC2-mediated transcriptional repression of germination-related metabolic genes is required to promote the seed-to-seedling developmental transition.

## Discussion

Seedling establishment involves an important developmental and metabolic transition. It is promoted by PRC2, which is evidenced by delayed seed germination (**Figure S5D**) (Bouyer et al., 2011; Müller et al., 2012) and a developmental reversal to an embryo-like state in PRC2-depleted plants (**Figure 4A and 5**) (Chanvivattana et al., 2004; Aichinger et al., 2009; Bouyer et al., 2011; Ikeuchi et al., 2015; Mozgová et al., 2017). However, how PRC2 reprograms the gene regulatory networks to prevent the developmental reversal and promote seedling establishment remained unknown. Here, we show that photosynthetic carbon assimilation is initiated between 3 - 5 days after germination initiation (dag) (**Figure 1A**). This coincides with the estimated depletion of storage reserves in Arabidopsis at 4 – 5 dag (Kircher and Schopfer, 2012; Xiong et al., 2013; Henninger et al., 2021) and supports a switch to photoautotrophy around the same time. In a recent study, first changes in H3K27me3 distribution in emerging seedling were observed only after 48 – 72 hrs after imbibition (Pan et al., 2023), which coincides with the onset of transcription of genes encoding PRC2 subunits (Mozgová et al., 2017; Pan et al., 2023). This timepoint seems to represent an important milestone in seedling establishment. Since reprogramming of H3K27me3 had not been studied beyond 72 hrs after imbibition (Pan et al., 2023), we focused on the transcriptome and H3K27me3 distribution in seedlings at 3 dag (heterotrophic) and 7 dag (photoautotrophic) plants, identifying two phases of PRC2-mediated gene repression (**Figure 6G**). First, in the still-heterotrophic seedling (before 3 dag), H3K27me3 is deposited and ensures irreversible repression of genes transcribed during seed development and embryo maturation, including the *LAFL* TF network and storage reserve biosynthesis genes. Second, between 3 and 7 dag, metabolic pathways related to germination, early seedling establishment are repressed.

Different TFs may target PRC2 recruitment to specific groups of genes during seedling establishment. PRC2 is recruited to cis-elements in the promoters of target genes by interacting with TFs (Xiao et al., 2017), including VAL1 and VAL2 (Yuan et al., 2021), TRB (Zhou et al., 2018), or AZF1 and BPC (Xiao et al., 2017). We show that H3K27me3 targets induced by external sucrose in *cs* (564 genes) and genes that gain H3K27me3 in photoautotrophic *cs* (126 genes) are exclusively enriched for VAL1/VAL2-targets (Yuan et al., 2021). Accordingly, the VAL1/VAL2-recognised RY-element is enriched among genes the gain H3K27me3 between 3 and 7 dag (**Figure S10B - D**). In contrast, TRB1 (Zhou et al., 2018) and AZF1/BPC1 (Xiao et al., 2017) targets and genes containing the REF6-binding motif (Pan et al., 2023) are enriched among genes losing H3K27me3 between 3 and 7-dag (378 genes). Mutation of REF6 did not affect H3K27me3 removal before 72 hrs after imbibition (Pan et al., 2023). Nevertheless, the presence of REF6-binding motifs in genes that lose H3K27me3 between 3 to 7-dag indicates a possible contribution of REF6 to the removal of H3K27me3 after 3 dag. Interestingly, the phenotype and transcriptome of sugar-grown mutants *val1 val2* resemble *clf swn* (Yuan et al., 2021), including the sugar-dependent accumulation of TAGs and activation of the *LAFLs* (Tsukagoshi et al., 2007; Suzuki et al., 2007; Yuan et al., 2021). In contrast, no TAG-accumulating phenotype has been reported in *trb1/2/3* combined mutants (Zhou et al., 2018; Wang et al., 2023) despite transcriptome similarity between *trb1/2/3* and *clf swn* (Zhou et al., 2018) and enrichment of TRB1 targets among H3K27me3-marked genes activated in photoautotrophic *cs* (805 genes, **Figure S10C**). These results suggest that developmentally separated modes of PRC2 recruitment exist. VAL1/VAL2-recruited PRC2 seems important for preventing the developmental reversal to embryonic state and for establishing vegetative seedling. In contrast, TRB1 and AZF1/BPC1 targets are enriched among genes losing H3K27me3 during seedling establishment, but TRB1 and/or AZF1/BPC1-mediated PRC2 recruitment can prevent gene activation in vegetative seedlings.

We show that the previously reported reactivation of the embryo maturation program (Chanvivattana et al., 2004; Aichinger et al., 2009; Bouyer et al., 2011; Ikeuchi et al., 2015; Mozgová et al., 2017) in PRC2-depleted seedlings is conditioned by external sucrose. In the absence of sucrose, or its addition after 3 dag, *LAFLs* transcription is stably repressed in *cs*, and no TAGs accumulate even without PRC2 (**Figure S5H**). Thus, the initial downregulation of the *LAFLs* and their downstream targets is independent of PRC2, but H3K27me3 deposition is required to prevent their reactivation in a narrow developmental time window, at which these genes can be potentially reactivated by external stimuli (e.g. sucrose). The observation is reminiscent of PRC2-mediated repression of the flowering repressor *FLC*, where PRC2 is required not to induce, but to maintain a cold-induced repressed state (Hepworth and Dean, 2015). This example further supports a role of PRC2 in stably “locking” a transcriptionally inactive state to prevent gene reactivation and developmental reversion.

In addition to promoting the developmental transition from seed to seedling, the timing of H3K27me3 deposition during seedling development may limit the potential of developmental reprogramming, that is utilized in biotechnological tissue and plant regeneration. Indeed, we found that within the first 7 days, H3K27me3 represses a number of *TFs* instructive for embryo, root or shoot development and regeneration (**Figure 6A, B**). Beside *LEC1* (Lotan et al., 1998) and *LEC2* (Stone et al., 2001), these include for example *WUS* (Zuo et al., 2002; Zhang et al., 2017), *CUC1/2* (Daimon et al., 2003), *AGL15* (Thakare et al., 2008), *PLT4/BBM* (Boutilier et al., 2002; Horstman et al., 2017), *PLT5/EMK* (Tsuwamoto et al., 2010; Kareem et al., 2015) or *PLT7* (Kareem et al., 2015). In agreement with H3K27me3 deposition at *LAFLs* before 3 dag, the potential to induce somatic embryo development, associated with *LAFL* reactivation (Jia et al., 2013; Horstman et al., 2017), is often restricted to seed germination or early seedling development (Horstman et al., 2017; Mozgová et al., 2017). Deposition of H3K27me3 during early seedling establishment is therefore likely to limit the time window of regenerative potential in plant tissues.

The deposition of H3K27me3 before 3 dag is independent of photosynthesis and light signaling (**Figure 6E**), and is mostly insensitive to the photosynthetic inhibitor DCMU (**Figure 3A**). The 3-dag seedlings were not yet assimilating photosynthetic sugar (**Figure 1A**), suggesting that they were photosynthetically inactive and therefore the impact of DCMU could be limited. In contrast, the levels of H3K27me3 dramatically decreased in the shoot and root of heterotrophic 7-dag plants (Figure 3A-C and Figure S3A, B). In this system, exogenous sucrose and DCMU was provided to light-grown plants to force heterotrophy. DCMU, that blocks electron transport from PSII to PQ_B_, also induces ROS production (Flors et al., 2006). Accordingly, we found that oxidative stress responses are activated in the DCMU-grown plants (**Table S2, S3**). Therefore, it is possible that the observed decrease of H3K27me3 and transcriptional response in 7-dag heterotrophic plants reflects a general oxidative stress response, rather than heterotrophic mode of nutrition and growth. Indeed, the distribution of H3K27me3 or transcriptome of 7-dag DCMU-grown (heterotrophic) shoot (S-H) did not resemble the 3-dag shoot (S-P3 or S-D3) (**Figure 1 C, D**), further corroborating this notion. We did not observe consistent changes in the transcription of PRC2 subunits or H3K27me3 demethylases that could directly explain the global and consistent effect of DCMU on H3K27me3 in both the shoots and the roots (**Figure S3C**). Thus, the mechanisms that led to H3K27me3 depletion in DCMU-treated sucrose-grown plants requires further investigation. Interestingly, transcriptional changes indicating elevated oxidative stress in photoautotrophic *cs* shoots resembled those observed in the shoot of DCMU-grown 7-dag plants (**Figure 4I**). Metabolic and transcriptomic changes in *cs* (**Figures 4, 5**) are also reminiscent of changes that occur in plants with reduced activity of TOR (Caldana et al., 2013). Strong inhibition of TOR inhibits nuclear PRC2 activity (Ye et al., 2022) and reduces global levels of H3K27me3 (Ye et al., 2022; Dong et al., 2023). It will be important to determine whether TOR signalling contributes to the DCMU-induced H3K27me3 depletion and how H3K27me3 depletion in *cs* feeds back into the potential crosstalk.

Primary metabolic genes are targeted by PRC2 during the transition to vegetative growth. These include important genes involved in lipid metabolism, the glyoxylate cycle or gluconeogenesis in the shoot, as well as photosynthesis-related genes in the root (**Figure 2**). Several observations in this work indicate that developmental identity can be underpinned by metabolic state. First, we found that inhibition of photosynthesis in the shoot is associated with the upregulation of photosynthesis-related genes and downregulation of root development in the root (**Table S3**). Second, the development of TAG-filled cotyledon-like structures in *cs* plants, instead of photosynthetically active true leaves, is promoted by the addition of exogenous sucrose (**Figure 4A** and **Figure S5E - H**). Third, preventing the transcription of *ICL* can promote vegetative transition in *cs* plants (**Figure 6F**). Although the molecular mechanisms are not yet clear, these results demonstrate a close link between the developmental and metabolic identities controlled by PRC2, and highlight the possibility of influencing developmental identity through metabolic interventions.

## Materials and methods

### Plant material and cultivation conditions

Wild type (WT) (Col-0) plants and strong PRC2 mutant alleles *clf-29* (SALK_021003) (Bouveret et al., 2006) and *swn-3* (SALK_050195) (Chanvivattana et al., 2004) were used in most experiments. For confirmation other strong PRC2 mutant alleles *clf-28 swn-7* (*clf-28:* SALK_139371; *swn-7*: SALK_109121) (Lafos et al., 2011; Gan et al., 2015), and *fie cdka;1* (*fie*: SALK_042962; *cdka;1*: SALK_106809) (Bouyer et al., 2011) were used. The combination of the used *clf* and *swn* alleles induce a severe developmental phenotype (Lafos et al., 2011; Mozgová et al., 2017), preventing homozygous seed production. Thus, *clf swn* (*cs*) double mutant plants were selected from a segregating population of seeds produced by *CLF/clf swn/swn* (*Ccss*) parental plants. The *cs* phenotype/genotype can be faithfully distinguished at 3 days after germination induction (dag). Parental generations of compared plants were grown concurrently under the same growth conditions.

The following cultivation conditions were used unless specified otherwise. The seeds were surface sterilised by ethanol and placed on one of the following media: “P” (photoautrophic): ½ MS (Murashige & Skoog medium: ½ Murashige & Skoog (MS) medium including vitamins (Duchefa cat.no. M0222), 2-(N-morpholino)ethanesulfonic acid (MES) and 0.8% (w/v) plant agar (Duchefa cat.no. P10011000)), “D” (DCMU): ½ MS + 10 μM DCMU (3-(3,4-dichlorophenyl)-1,1-dimethylurea: Diuron – Sigma Aldrich cat.no. D2425), “M” (mixotrophic): ½ MS + 30mM (1%) sucrose; “H” (hetrotrophic): ½ MS + 30mM (1%) sucrose + 10 μM DCMU. For ^13^C analyses, MS medium without vitamins was used (Duchefa cat.no. M0221). Sterile layer of cellophane was placed on the top of the solidified medium to avoid medium-origin carbon contamination. For selected experiments, 30 mM mannitol was used as osmotic control. The seeds were stratified over 2 nights (ca 65 hrs) and placed in long-day conditions (110-120 µmol m^-2^ s^-1^, 16hrs light/8 hrs dark, 22 °C during light/20 °C during dark; light source: OSRAM FQ 54W/840 HO CONSTANT LUMILUX Cool White). Germination (radicle emergence) was scored at given time points, presence of *clf swn* phenotype in the population of *CLF/clf swn/swn* progeny was scored at 10 days after germination induction (dag). Plant in soil for seed amplification and scoring seed set and abortion rate were cultivated under long-day conditions (110 µmol m^-2^ s^-1^, 16hrs light/8 hrs dark, 21 °C).

### Generation of CRISPR-ICL lines

To target *ISOCITRATE LYASE (ICL)* (AT3G21720), two suitable sgRNA targeting *ICL* - *exon 2* were designed using CRISPOR (Concordet and Haeussler, 2018) and cloned in pDGE Shuttle vectors (Addgene plasmid #153241 pDGE332 and Addgene plasmid #153243 pDGE334) before assembly into pDGE347 (Addgene) using the method previously described (Stuttmann et al., 2021). The binary vector was transferred to *Agrobacterium tumefaciens* (GV3101) for transformation by floral dipping (Clough and Bent, 1998) of *CLF/clf-29 swn-3/swn3* (*Ccss*) plants. All sgRNAs and oligonucleotides are listed in (Table S25). RFP(Cas9)-positive T1 seeds were selected using Leica S9i (Leica, DE) stereomicroscope with NIGHTSEA fluorescence adaptor (Nightsea, USA) and cultivated in soil. *Ccss* plants were selected and *ICL* locus containing the targeted sites was sequenced by Sanger sequencing (Table S25). RFP(Cas9)-negative T2 seeds were selected and *Ccss icl* homozygous mutant plants were selected. 10 independent *Ccss icl* lines were analysed that showed comparable phenotypes, 4 lines are shown.

### Detection of TAGs using Sudan Red 7B (fat red)

21-dag plants were analysed for the presence of embryonic lipids using Sudan Red 7B (Sigma-Aldrich, #46290) as described by Aichinger et al. (Aichinger et al., 2009). Briefly, seedlings were immersed in filtered solution of 0.5% (w/v) Sudan Red 7B in 60% isopropanol for 15 minutes and rinsed three times with H_2_O. The penetrance of fat-red phenotype was calculated as percentage of red-staining plants. Phenotypes were documented using Optika stereo-microscope equipped with optikam PRO8 Digital Camera (Optica, IT) or Leica S9i stereomicroscope (Leica, DE).

### Determination of CO_2_ assimilation

Two approaches were taken to assess CO_2_ assimilation. To assess the onset of CO_2_ assimilation (**Figure 1A**), we used natural ^13^C fractionation, exploiting the different isotopic composition of CO_2_ in atmospheric air (δ^13^C = -8.5‰) and artificial air (δ^13^C = -40‰). Stratified seeds of plants grown in atmospheric air (δ^13^C seeds ∼ -30‰, reflecting metabolic ^13^C discrimination (Cernusak et al., 2013)) were placed into a desiccator continually flushed with artificial air (mixture of ∼80% N_2_, 20% O_2_ and 0.04% CO_2_; δ^13^C = -40‰) and plants were grown for up to 14 dag. Decrease in δ^13^C in 14 dag plants results in δ^13^C_shoot_ ∼ -55‰ and δ^13^C_root_ ∼ - 50‰, reflecting the ^13^C signal of assimilated CO_2_ in artificial air, ^13^C discrimination during photosynthesis (Farquhar et al., 1989) and post-photosynthetic discrimination (Cernusak et al., 2009). To assess the level of CO_2_ assimilation in 1 hr (**Figure 4B**), a ^13^C-labelling approach was carried out as previously described by Kubásek et al. (Kubásek et al., 2021). In particular, seedlings were cultivated under standard long-day conditions (as specified above) for 10-days and ^13^C labelling was carried out at ZT 7 – 8 (middle of photoperiod). Open plates with seedlings were placed in a 3-litre chamber made of an inverted glass petri dish with water at the bottom to seal the gap between the dish and the lid against gas leakage. The chamber was equipped with a sealed metal tube to allow gas injection and a ventilator for homogeneous gas distribution. The chamber was flushed with 5 l of CO_2_-free air (80% nitrogen, 20% oxygen) to remove ambient CO_2_. Next, 3 ml of ^13^CO_2_ (99% ^13^C, 0% ^12^C) were injected into the chamber (final CO_2_ concentration was 1000 µmol.mol^-1^), after which the plants were incubated in the closed chamber for 1 hour (pulse). For both approaches, shoots and roots were dissected and air-dried to obtain final 0.1 – 0.2 mg DW material per sample. Samples were combusted in oxygen (elemental analyser Flash 2000, Thermo Scientific, Brehmen, Germany) and the ^13^C/^12^C ratio in the resultant CO_2_ was determined by isotope ratio mass spectrometer (Deltaplus XL, Thermo Scientific, Brehmen, Germany). Each experiment was performed in biological triplicates.

### Biomass analysis

WT (Col-0) and *clf-29 swn-3* seeds were sterilised and plated on ½ MS medium plates “P” (photoautrophic) and “M” (mixotrophic) and cultivated under standard long-day conditions (as previously described). For the biomass analysis WT and *cs* seedling were sampled at 6 dag and 12 dag. The fresh weigh of the seedlings was assessed for the biomass analysis. The experiment was performed using four biological replicates.

### ABA concentration measurements

Extraction and purification of ABA was done using a previously described method (Turečková et al., 2024) with minor modifications. Frozen samples (5 mg fresh weight) were homogenized using a MixerMill (Retsch GmbH, Haan, Germany) and extracted in 1 ml of ice-cold extraction solvent methanol/water/acetic acid (10/89/1, v/v/v) and stable isotope-labelled internal standard (5 pmol of ^6^H_2_- ABA per sample added). The extracts were purified on Oasis HLB columns (30 mg, Waters Corp., Milford, USA), conditioned with 1 mL methanol and equilibrated with 1 mL methanol/water/acetic acid (10/89/1, v/v/v). After sample application, the column was washed with 1-mL methanol/water/acetic acid (10/89/1, v/v/v) and then eluted with 2 mL methanol/water/acetic acid (80/19/1, v/v/v). Eluates were evaporated to dryness and dissolved in 30 ul of mobile phase prior to mass analysis using an Acquity UPLC^®^ System and triple quadrupole mass spectrometer Xevo TQ MS (Waters, Milford, MA, USA) (Floková et al., 2014).

### Plant material for transcriptome, H3K27me3 and metabolome profiling

WT (Col-0) and *clf-29 swn-3* (*cs*) were used for all profiling experiments. Plants for profiling were cultivated within the same period of time and material harvested at the same time was divided to be used in all profiling experiments. Biological replicates were cultivated on separate plates, that were regularly moved within the cultivation chamber to randomize the environmental effects. For RNA-seq and ChIP-seq, WT plants were harvested at 3 and 7 dag. Due to 2-to 3-day delay in the germination, *cs* plants were harvested at 9-dag (growth shifted forward, harvesting done at the same time as 7-dag WT). Separated shoots and roots were collected by seedling dissection, and the same seedling pool was used as the corresponding shoot and root sample for RNA-seq and ChIP-seq. In 3-dag samples, material from 200 or 400 seedlings per replicate was pooled for RNA- or ChIP-seq, respectively (material from 400 seedlings corresponded to ca 50 mg FW for either tissue). In 7-dag WT, material from 100 or 200 seedlings per replicate was pooled for RNA- or ChIP-seq, respectively, and 50 mg FW was used in ChIP. In 9-dag *cs* samples, material from 150-200 seedlings per replicate was pooled for RNA extraction. Replicates A, B, and C were harvested at zeitgeber time (ZT) 4 – 7, ZT 7 – 10 and ZT 10 – 13, respectively. Each individual sample was collected into RNAlater (Thermo, ct. no. AM7020) and flash frozen in liquid nitrogen for RNA extraction or immediately crosslinked and flash frozen for ChIP. All replicates were used for RNA-seq and but only replicates B and C used in ChIP-seq, while replicate A was retained for qPCR confirmation. For primary metabolome profiling, five biological replicates of 20 mg FW shoot tissue per replicate were dissected from 7-dag WT and 9-dag *cs* seedlings. Collection was performed at ZT 4 – 6. Following shoot collection, the tissue was washed by MilliQ and dried on filter paper before flash freezing in liquid nitrogen.

Seed stages for metabolome profiling were harvested from the main inflorescence of WT (Col-0) plants grown on soil. Siliques were tagged at pollination (representing stage 1) and harvested in pools of 2-3 successive siliques per stage, obtaining 25 stages along the main inflorescence stem (stage 25 representing the most mature siliques in the bottom part of the inflorescence stem). In stages 6 – 25, developing seeds were dissected from the lyophilised siliques. Earlier stages were combinations of siliques and developing seeds. Samples were pools of 3 mg FW, representing a pool of 170-200 plants each. Three independent replicates were harvested at each day after pollination, day 22 had two replicates, days 24 and 25 yielded only a single replicate.

### Gene transcription quantification by RT-qPCR

WT (Col-0) and *clf-29 swn-3* seeds were sterilised and plated on ½ MS medium plates “P” (photoautrophic) and “M” (mixotrophic) and cultivated under standard long-day growth conditions (as previously described). WT seedlings were harvested at 2 dag (∼ 200 seedlings per replicate) and 7 dag (∼100 seedlings per replicate), and *cs* seedlings were harvested at 9 dag (∼ 200 seedling per replicate) at ZT 4 – 7. Each sample were collected separately into RNAlater (Thermo, ct. no. AM7020) and immediately flash frozen in liquid nitrogen for RNA extraction. 50 mg of collected material from each sample were used for the RNA extraction. RNA was extracted using MagMAX™ Plant RNA Isolation Kit (Thermo, cat.no. A33784). 1 μg of extracted RNA was reverse-transcribed using the RevertAid First Strand cDNA Synthesis Kit (Thermo, cat.no. K1622) with Oligo (dT)_18_ primers. RT-qPCR was performed using the CFX Connect^TM^ Optics Module Real Time PCR detection system (BIO-RAD) with gene-specific primers (Table S25) and 5x HOT FIREPol Eva Green qPCR Mix Plus (ROX) (Solis Biodyne, cat.no. 08-25-00008). The experiment was performed using three biological replicates. *PP2A* (*AT1G13320*) was used as reference gene. R programming language version R 4.3.1 (R Core Team, 2023) was used for statistical computing and graphics. Outliers were excluding using Dixon’s (*p*-value threshold = 0.05) tests with the R package Outliers (Komsta, 2022). Two-way ANOVA tests were performed with base R and Tukey’s HSD test tests were performed with emmeans (Lenth, 2016) and mulcomp (Hothorn et al., 2008) and heatmaps were produced with pheatmap (Kolde, 2019), RColorBrewer (Neuwirth, 2022), ggplotify (Yu, 2019), and ggplot2 (Hadley Wickham, 2016) R packages. Hierarchical clustering was based on row Z-score-normalized transcript abundance of genes in 2- and 7-dag photoautotrophic (S-P2 and S-P) and mixotrophic (S-M2 and S-M) WT shoot and 9- dag photoautotrophic (*cs*-S-P) and mixotrophic (*cs*-S-M) *cs* shoot.

### RNA-seq library preparation, sequencing and data analyses

Total RNA was extracted from root and shoot tissues using MagMAX™ Plant RNA Isolation Kit (Thermo, cat.no. A33784). polyA-mRNA was enriched using NEBNext® Poly(A) mRNA Magnetic Isolation Module (NEB, cat. No. E7490) and Illumina sequencing libraries were prepared using NEBNext® Ultra™ II RNA Library Prep Kit for Illumina® (NEB, cat. No. E7770), 24 libraries were pooled and sequenced on Illumina HiSeq4000 in 50 bp single-end (SE) mode. Obtained numbers of reads per sample are available in (Table S1). Raw RNA-seq data were trimmed using TrimGalore! v0.4.1 (https://github.com/FelixKrueger/TrimGalore, v0.4.0)., quality checked using FastQC v0.11.5 (Andrews, 2010) and mapped to Arabidopsis thaliana TAIR10 genome using Hisat2 v2.0.5 (Kim et al., 2019). Gene expression was quantified as RPKM (reads per kilobase of the gene per million of the reads in the dataset) in Seqmonk v1.40.0 (https://www.bioinformatics.babraham.ac.uk/projects/) using Araport11 annotation. Differential gene expression was analysed using DESeq2 and EdgeR within Seqmonk v1.40.0 with *p*-value cut-off 0.05. Genes were considered significantly differentially expressed if they were identified as such by both DESeq2 and EdgeR and the expression fold change was at least +/- 1.5 (abs. log_2_FC 0.6).

### Chromatin immunoprecipitation and qPCR

Chromatin immunoprecipitation was carried out using previously described protocol (Mozgová et al., 2015). In brief plant material was crosslinked using 1% formaldehyde for 10 min under vacuum and crosslinking was terminated using 0.125 M glycine for 5 min. Sheared chromatin (10 cycles, 30 s ON/ 30 s OFF using a Bioruptor® (Diagenode)) extracted from 100 mg material was equally divided between input and samples immunoprecipitated with anti-histone H3 (Merc, cat. no. 07-690); anti-H3K27me3 (Merc, cat. no. 07-449) and IgG (Merc, cat. no. I5006) as control. Immunocomplexes were collected using Dynabeads® Protein A for Immunoprecipitation (Thermo, cat. no. 10001D) and washed 2 x 5 min with low-salt buffer (containing 150 mM NaCl), 2 x 5 min with high-salt buffer (containing 500 mM NaCl) and 1 x 5 min with LiCl-containing buffer. Recovered DNA was extracted using phenol-chloroform and precipitated by ethanol with addition of GlycoBlue™ Coprecipitant (Thermo, cat. no. AM9515) and purified DNA was analyzed by qPCR using CFX Connect™ Real-Time PCR Detection System (Bio-Rad) using 5X HOT FIREPol Eva Green qPCR Mix Plus (ROX) (Solis Biodyne) with gene-specific primers detailed in (Table S25). ChIP performance was quantified by comparing immunoprecipitated DNA to input and abundance of H3K27me3 is shown as anti-H3K27me3 or IgG related to anti-H3 recoveries (H3K27me3/H3 or IgG/H3).

### ChIP-seq library preparation, sequencing and data analyses

For ChIP-seq, immunoprecipitated DNA was purified using iPure kit v2 (Diagenode, cat. no. C03010015). Following qPCR control to assess locus-specific signal/noise ratio, DNA amount in input, anti-H3 and anti-H3K27me3 immunoprecipitated was quantified using Qubit® dsDNA HS Assay (Thermo, cat. no. Q32854) and 3 ng of DNA was used for Illumina sequencing library preparation using NEBNext® Ultra™ II DNA Library Prep Kit for Illumina® NEB, cat. no. E7645S) according to manufactureŕs instructions. Libraries were pooled into two pools of 24 samples (expected output min 20 Mio reads per sample) and sequenced on the Illumina HiSeq Hi-Outputv4 platform in 125 bp paired-end (PE) mode. Obtained numbers of reads per sample are available in (Table S4). To assess the quality of the sequence (raw and after trimming), FastQC v0.11.5 (Andrews, 2010) was used. To clean the raw data, TrimGalore v0.6.2 was used, with parameters: Paired; Trim N; Phred33; Stringency 6; Quality 20; Minimum length 20; Clip 10 bp from 3’ and 5bp from 5’. Raw ChIp-Seq reads were mapped to TAIR10 *Arabidopsis thaliana* genome using Bowtie2 v2.2.9 enabling the option --no-mixed. The SAM file was processed (mapping quality filter, with a MAPQ threshold of 25, conversion to BAM format, sorting and indexing) using SAM tools v1.9. Following Spearman correlation analyses of the two biological replicates, reads from the two replicates were pooled in the final analyses. Coverage of input, H3 and H3K27me3 was computed using deepTools2 – BAMcoverage (Ramírez et al., 2016). Reads were extended to 200 bp (estimated mean library insert size), normalized by BPM (bins per million mapped reads), bin size was 50bp. Enrichment of H3K27me3/H3 was computed per 50 bp bin (in log_2_ ratio) with deepTools2 – BAMcompare taking H3K27me3 as sample and H3 as control (Ramírez et al., 2016). Average enrichment per feature (annotated gene) was calculated using DeepTools2 – multiBigwigSummary (Ramírez et al., 2016), using H3K27me3/H3 enrichment values. BedTools v2.26.0 was used to parse the BAM format to BED format. SICER was used to identify enriched regions (peaks) for H3K27me3, broad modification, with the following parameters: Redundancy threshold 25; Window size 200; Gap 600; Effective genome size 0.998444; FDR 0.05; Fragment size 150; Control: H3 enriched sample; Sample: H3K27 enriched sample. To relate features (genes) to identified enrichment peaks, BedTools v2.26.0 (Intersect) and Araport11 annotation was used (Cheng et al., 2017). To identify a feature as covered by a peak, at least 1 bp of the feature must be covered by the peak (T1 genes), or ≥70% of feature body must be covered by the peak (T70 genes). A peak could cover more than one feature.

### Gene ontology enrichment and TF gene identification

Gene ontology enrichment analysis was performed using DAVID 6.8 (Langmead and Salzberg, 2012). GO categories passing the thresholds of Bonferroni-corrected *p*-value ≤ 0.05 and min. enrichment ≥ 1.5 were considered. Bubble plots of GO-enriched pathways were plotted using SRplot (Tang et al., 2023). TF-gene association was based on Plant Transcription Factor Database (PlantTFDB - https://planttfdb.gao-lab.org/) and TF genes defined by Czechowski et al. (Czechowski et al., 2004).

### Enrichment of VAL1/VAL2, TRB1 and AZF1/BPC1 targets, and PRE motif enrichment

Lists of gene targets of VAL1/VAL2 (Yuan et al., 2021) and TRB1 (Zhou et al., 2018) were taken from the respective studies. AZF1 and/or BPC1 targets were determined from the centre of ChIP-peak coordinates (Xiao et al., 2017) using ChIPSeeker R package (Wang et al., 2022). Peaks centred within 3kb upstream and 250bp downstream of TSS of Araport11-annotated gene were assigned to that gene. In case of multiple assignments, the gene whose TSS was the closest was retained. 9036 genes marked by H3K27me3 in 3- and/or 7-dag shoot of WT seedlings were used as background. Polycomb response element (PRE) and REF6-binding motif enrichment was analysed in promoter sequences (-1000 to 0 bp from TSS), retrieved by RSAT (TAIR10 assembly) (Nguyen et al., 2018). Position weight matrices were obtained from publications by Zhou et al. (Zhou et al., 2018) and Yuan et al. (Yuan et al., 2021). Promoter sequences of the whole genome were used as a background and the p-values were computed with the sea (simple enrichment analysis) command of the meme suite (Bailey et al., 2015), considering a 3-order model (options: --order 3 --align right --hofract 0.3).

### Metabolite extraction and data analysis

Deep frozen plant shoot samples of ∼20 mg fresh weight (FW) or ∼ 3 mg (FW) of developing seed material were homogenized in the frozen state by an oscillating ball mill, extracted by pre-cooled solvents, chemically derivatized, and analyzed by gas chromatography - electron impact ionization - mass spectrometry (GC-MS) as described previously by Erban et al. (Erban et al., 2020). In detail, 160 µL polar phase of an extraction mixture of 360:400:200 (v/v/v) methanol:water:chloroform were obtained after thorough mixing with the frozen powder and separation of liquid phases by centrifugation for 5 min at 20,800 x g. ^13^C_6_-Sorbitol was added as internal standard to the extraction mixture. The 160 µL aliquot was dried in a vacuum concentrator and stored at -20 °C until further processing. In the case of seedling analysis, the complete polar phase was sampled and dried. Before GC-MS injection, dry samples were methoxyaminated by methoxyamine hydrochloride dissolved in pyridine to a final concentration of 40 mg * mL^-1^ and subsequently trimethylsilylated by N,O-bis(trimethylsilyl)trifluoroacetamide (Erban et al., 2020). A standard mixture of *n*-alkanes, *n*-decane (RI 1000), *n*-dodecane (RI 1200), *n*-pentadecane (RI 1500), *n*-octadecane (RI 1800), *n*-nonadecane (RI 1900), *n*-docosane (RI 2200), *n*-octacosane (RI 2800), *n*-dotriacontane (RI 3200), and *n*- hexatriacontane (RI 3600), was prepared in pyridine and added to each derivatized sample for retention index calibration. Metabolite profiles were performed by an Agilent 6890N gas chromatograph with split/splitless injection and electronic pressure control (Agilent, Böblingen, Germany) using a 5% phenyl–95% dimethylpolysiloxane fused silica capillary column of 30 m length, 0.25 mm inner diameter, 0.25 μm film thickness, an integrated 10 m pre-column and helium carrier gas for chromatographic separation. Electron impact ionization and time-of-flight mass spectrometry were performed with a Pegasus III time-of-flight mass spectrometer (LECO Instrumente GmbH, Mönchengladbach, Germany) (Erban et al., 2020).

ChromaTOF software (Version 4.22; LECO, St. Joseph, USA) and TagFinder software (Luedemann et al., 2008) were used for chromatogram processing and compound annotation. Analytes, i.e. chemically derivatized metabolites, were annotated manually using TagFinder software by matching of mass spectral and chromatographic retention index information to reference spectra and retention indices of authenticated reference compound from the Golm Metabolome Database (http://gmd.mpimp-golm.mpg.de), metabolite, analyte, and match quality information are reported (Tables S17-S18). Each sample was analyzed in both, splitless and split (1:30 ratio) mode. Abundant compounds that were above the upper limit of quantification in splitless mode were re-analyzed by split mode. Acquired metabolite abundances of arbitrary units were background-subtracted by non-sample control measurements, normalized to the abundance of the internal standard ^13^C_6_-sorbitol and to the sample fresh weight of each sample for relative quantification by normalized metabolite responses, arbitrary units g^-1^ (FW).

R programming language version R 4.3.1 (R Core Team, 2023) was used for statistical computing and graphics. Outliers were excluding using Rosner’s generalized extreme studentized deviate procedure (maximum number of outliers = 2, p-value threshold = 0.05) with PMMCRplus (Pohlert, 2023). Two-way ANOVA with Tukey’s HSD test and heatmaps were produced using R packages as described in previous section. Hierarchical clustering is based on row Z-score-normalized relative abundances of 85 metabolites in WT photoautotrophic (S-P) and mixotrophic (S-M) shoot and *cs* photoautotrophic (cs-S-P) and mixotrophic (cs-S-M) shoot samples. Principal Component Analysis (PCA) of metabolite profiles (Table S17) was performed in R 4.1.1 (R Core Team, 2021). Normalized response data were log_10_-transformed, mean-centered and auto-scaled prior to PCA and processed using the packages ggplot2 (Hadley Wickham, 2016), ggpubr (Kassambara, 2020), and factoextra (Kassambara and Mundt, 2021).

WT and *cs* mutant profiles were compared to time series of *Arabidopsis thaliana* seed maturation (Table S18) and germination data (Ginsawaeng et al., 2021) by Spearman rank correlation across the shared metabolites among the data sets (Table S21). Metabolite data were maximum-scaled per metabolite and dataset to a range of 0-100. All data were from GC-MS profiling analyses except amino acid abundances of the germination series that were LC-MS measurements (Ginsawaeng et al., 2021). Correlation coefficients of all sample combinations were arrayed in a symmetric matrix, tested for the significance of correlation (*P* > 0.05) and visualized as a three-color heat map in the range of negatively correlated (-1) in blue, non-correlated (0) in white, and positively correlated (+1) in red. Matrices of Spearman rank correlation coefficients from primary metabolome profiles of sucrose-supplemented WT, mannitol-supplemented mutant and sucrose-supplemented mutant correlated to the profiles of seed-maturation and seed-germination stages differed from mannitol-supplemented WT in a characteristic developmental stage dependent manner (Table S21). These differences of correlation coefficient matrices were tested at each developmental stage by the heteroscedastic Student’s t-test of the MS-EXCEL table calculation program and confirmed by Wilcoxon rank sum testing using the R function wilcox.test of the stats package (R Core Team, 2021). The correlation coefficient matrices were generated from correlation calculations of all available replicate measurements, namely 3-5 replicate profiles per seed maturation stage from this study or 5 replicates of the seed germination stages (Ginsawaeng et al., 2021) correlated to 6 replicates of the WT and mutant seedlings of this study, respectively. With few exceptions, normality of the correlation coefficient matrices, treatment groups versus developmental stages, was confirmed by Shapiro-Wilk testing, as well as heteroscedasticity by the Levene’s test (p ≤ 0.05).

Modules (=clusters) and metamodules (=metaclusters) of metabolites (**Figure S9**) were obtained using mean values maximum-scaled per metabolite and dataset. Weighted metabolites correlation network analysis implemented with the WGCNA R package (Langfelder and Horvath, 2008) was performed. Weighted networks rely on metabolite adjacencies, which correspond to absolute metabolite correlations exponents of a soft-thresholding power β. The value β = 7 was chosen to obtain a network with Scale-Free Topology properties (Scale Free Toplogy Model Fit R2 ≥ 0.6) and high connectivity.

Adjacency values were used to compute the Topological Overlap Matrix (TOM). Metabolites were clustered into modules based on the (1-TOM) distance matrix, by hierarchical clustering with average linkage distance comparison between clusters. The number of modules was determined by Dynamic Hybrid tree cut (dendrogram cut height for module merging = 0.25). Metamodules were identified based on the results of hierarchical clustering of the eigenmetabolites representing each module, considering the (1-TOM) distance.

### Statistical analyses and plotting

Statistical analysis and graphics generation were performed using OriginPro 2023, version 10.0.0.154 (Academic) and R version R 4.1.1 (R Core Team, 2021) and version R 4.3.1 (R Core Team, 2023).

## Data availability

The data for this study have been deposited in the European Nucleotide Archive (ENA) at EMBL-EBI under accession number PRJEB80531 (https://www.ebi.ac.uk/ena/browser/view/PRJEB80531).

## Author contributions

**NS**: experimental work including transgenic plant generation, data visualization, interpretation, manuscript writing and revision; **MTA, LG**: NGS data analyses; **QR**: metabolic cluster analyses; data visualization; cis-element analysis; **FA, HHM, MZ**: experimental work (sample collection, individual RT/ChIP-qPCR experiments); **AEr, AEb, AS, JK**: primary metabolome analyses, visualization, interpretation; **JK, JS**: CO_2_ assimilation experiments and data interpretation; **ON**: ABA content analysis; **AD, DB, FR**: CRISPR/Cas9 design and guidance; **IM:** conceptualization, funding, experimental work, draft writing and revision.

## Acknowledgements

Computational resources were provided by the e-INFRA CZ project (ID:90254), supported by the Ministry of Education, Youth and Sports of the Czech Republic. pDGE plasmids were a kind gift from Johannes Stuttmann and Sylvestre Marillonnet. We would like to thank Tomáš Konečný and Marie Vitásková for technical assistance.

## Supplemental Figures

**Figure S1.**
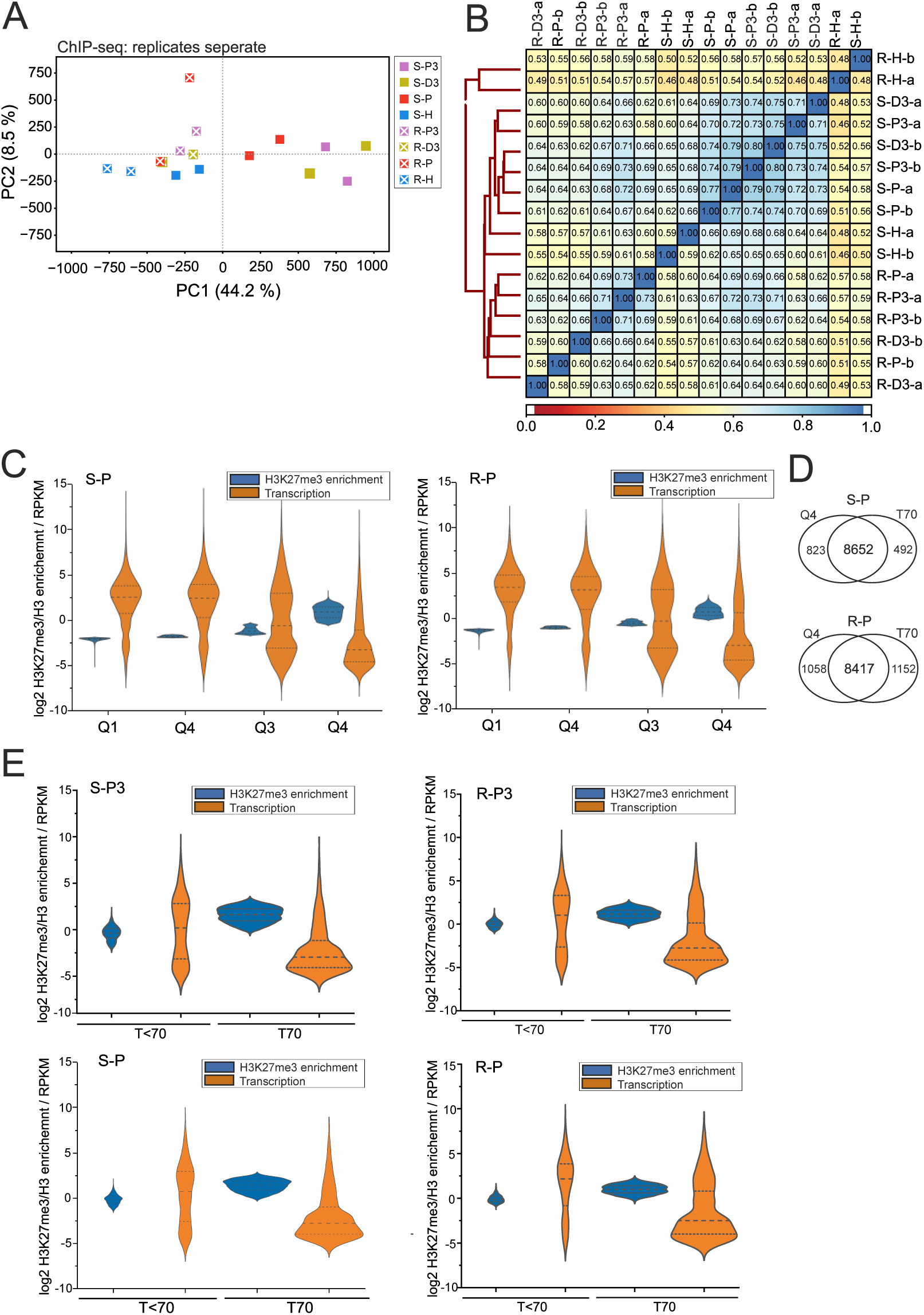
ChIP-seq technical controls: High level of H3K27me3 enrichment associates with low transcription. (Related to Figure 1). **A)** ChIP-seq principal component analysis (PCA) plot: genome-wide H3K27me3/H3 values in 200-bp windows, separate biological duplicates. **B)** Spearman correlation of analysed ChIP-seq samples. For final analyses, the reads of the respective biological duplicates were merged. **C)** Association between H3K27me3/H3 enrichment and gene transcription (RPKM). All Araport11-annotated genes were separated into quartiles (Q1 – Q4) based on the level of H3K27me3/H3 enrichment. **D)** Overlap between genes identified as most-highly enriched for H3K27me3 (Q4) and genes with ≥70% gene body overlapping with H3K27me3/H3-enriched region (T70). **E)** Association between H3K27me3/H3 enrichment and gene transcription (RPKM) in photoautotrophic 3- and 7-dag shoot and root tissues. T<70: genes with >0%<70% gene body or T70: genes with ≥70% gene body overlapping with H3K27me3/H3-enriched region. 3-dag photoautotrophic shoot (S-P3); 7-dag photoautotrophic shoot (S-P); 3-dag photoautotrophic root (R-P3); 7-dag photoautotrophic root (R-P). Similar patterns were observed for all samples.

**Figure S2.**
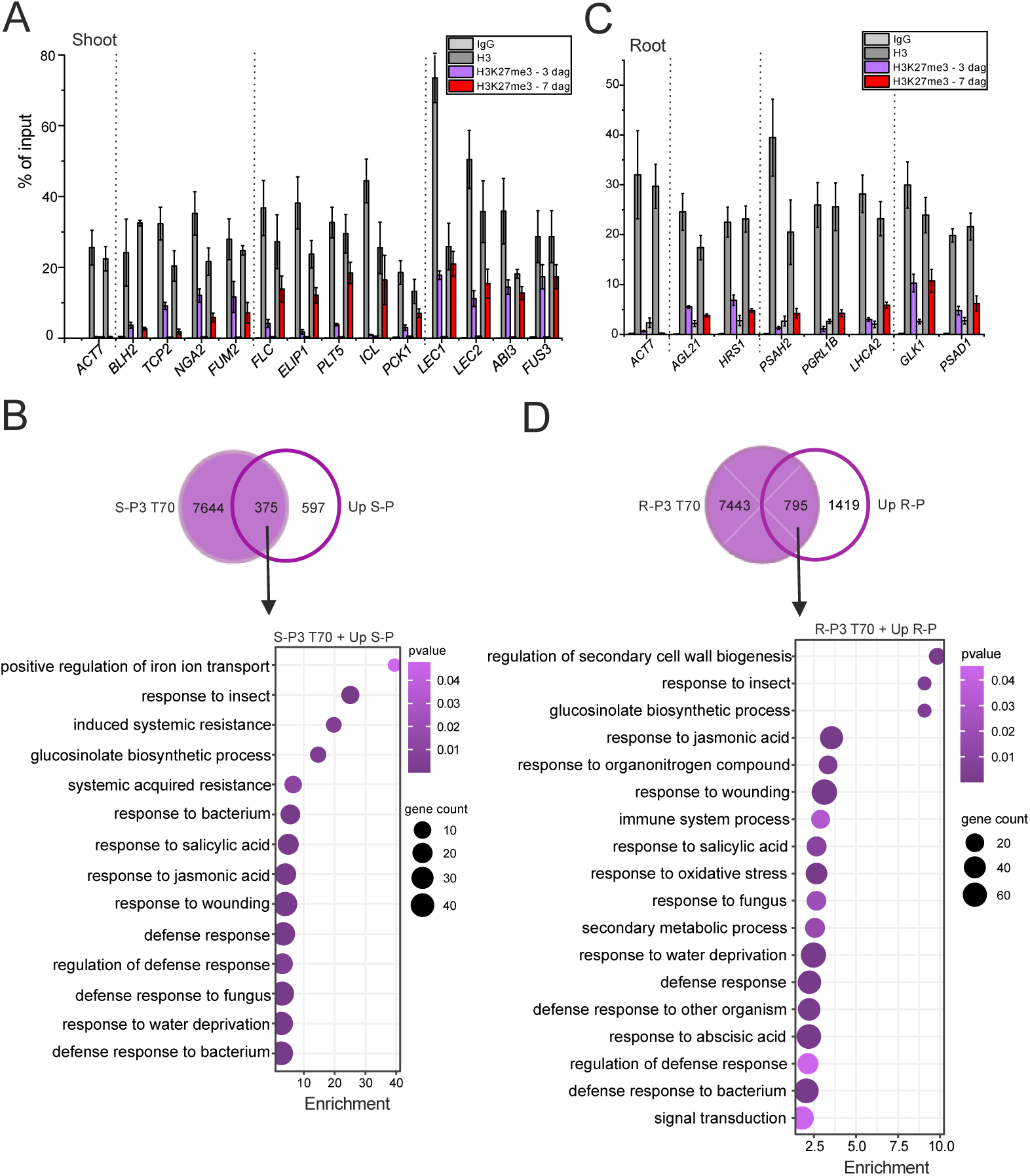
ChIP-qPCR confirmation of ChIP-seq and GO of genes that lose H3K27me3 and are activated between 3 and 7 dag. (Related to Figure 2). **A)** Full version of Figure 2C including ChIP controls: ChIP-qPCR analysis of representative genes with decreased (*BLH2, TCP2, NGA2* and *FUM2*), increased (*FLC, ELIP1, PLT5, ICL* and *PCK1*) or unchanged (*LEC1, LEC2, ABI3* and *FUS3*) levels of H3K27me3 from 3- to 7-dag shoot. *ACT7* serves as negative control locus with no H3K27me3 enrichment. Bars: mean ±SD; N = 3 technical replicates. **B)** Gene ontology (GO) enrichment of genes that lose H3K27me3 and are transcriptionally upregulated between 3- and 7-dag shoot. S-P3: 3-dag shoot; S-P: 7-dag shoot; T70: H3K27me3 target genes. BP categories are shown; GO display cutoff: fold enrichment > 1.5; p(Bonferroni) < 0.05. **C)** Full version of Figure. 2H with ChIP controls: ChIP-qPCR analysis of representative genes showed decreased (*AGL21* and *HRS1*), increased (*PSAH2* and *PGRL1B*), and unchanged (*LHCA2, GLK1* and *PSAD1*) levels of H3K27me3 from 3- to 7-dag root. *ACT7* serves as negative control locus with no H3K27me3 enrichment. Bars: mean ±SD; N = 3 technical replicates. **D)** GO enrichment of genes that lose H3K27me3 and are transcriptionally upregulated between 3- and 7-dag root. R-P3: 3-dag root; R-P: 7-dag root; T70: H3K27me3 target genes. BP categories are shown; GO display cutoff: fold enrichment > 1.5; p(Bonferroni) < 0.05.

**Figure S3.**
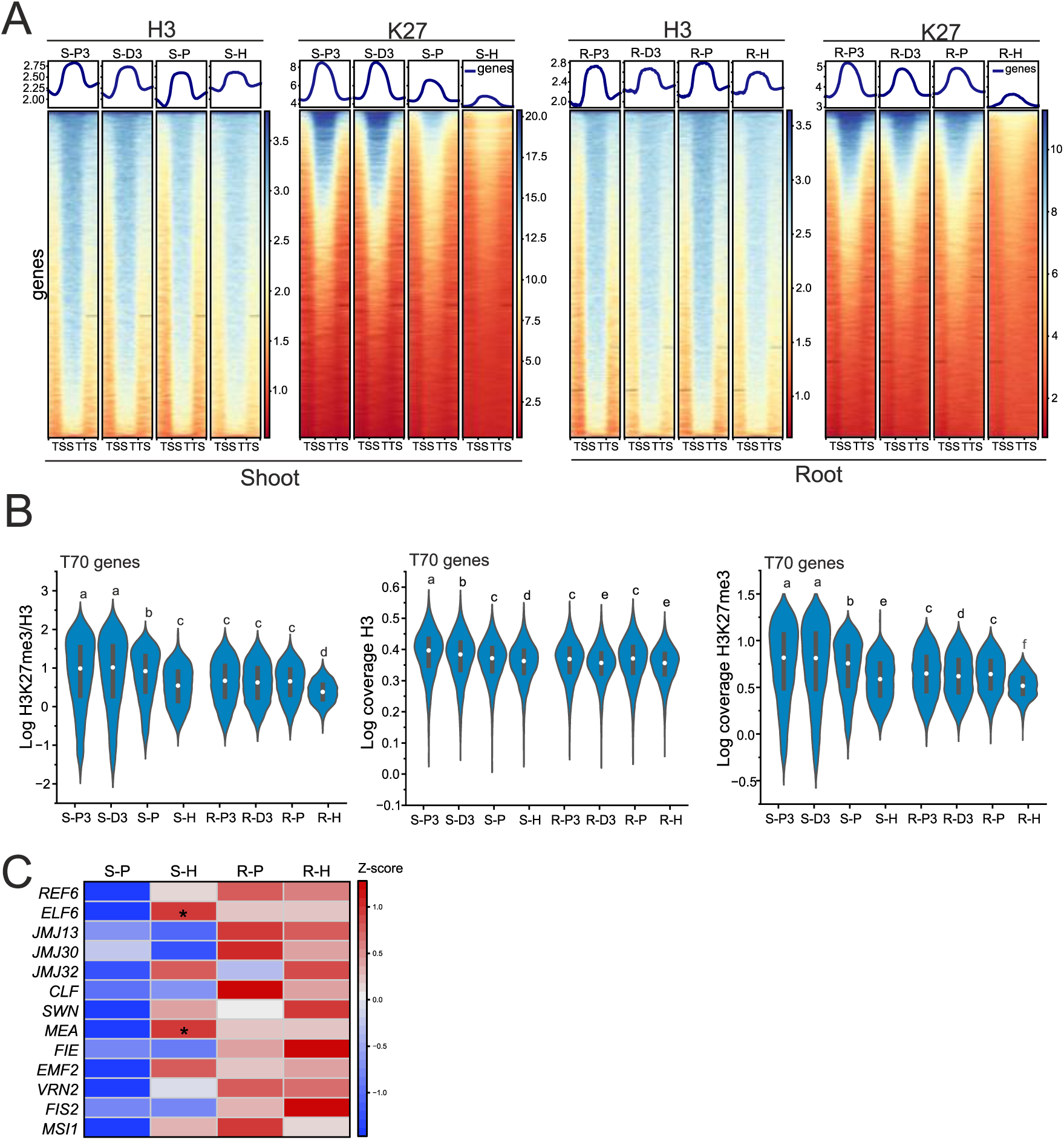
Coverage of H3 and H3K27me3 in H3K27me3-target genes identified in the shoot or root tissues. (Related to Figure 3). **A)** Coverage of H3 and H3K27me3 in gene bodies (-/+ 0.6 kb) in shoot and root samples for Araport 11–annotated genes. **B)** Enrichment of H3K27me3/H3 or coverage of H3 and H3K27me3 in genes identified as H3K27me3-targets (T70) in at least one of the profiled samples. Letters above the violin distribution plots: significance levels, p < 0.01: Dunńs multiple comparison test following Kruskal-Wallis ANOVA. Sample labels in A) and B) correspond to Figure 1B. **C)** Relative transcription of JMJ-domain H3K27me3 demethylase genes and PRC2 subunit genes in 7-dag photoautotrophic (P) or heterotrophic (H) shoot (S) or root (R). Heatmap represents Z-score normalized mean RPKM values. Asterisks (*): significant difference between respective (P) and (H) samples; p < 0.05: DESeq2; N = 3 biological replicates.

**Figure S4.**
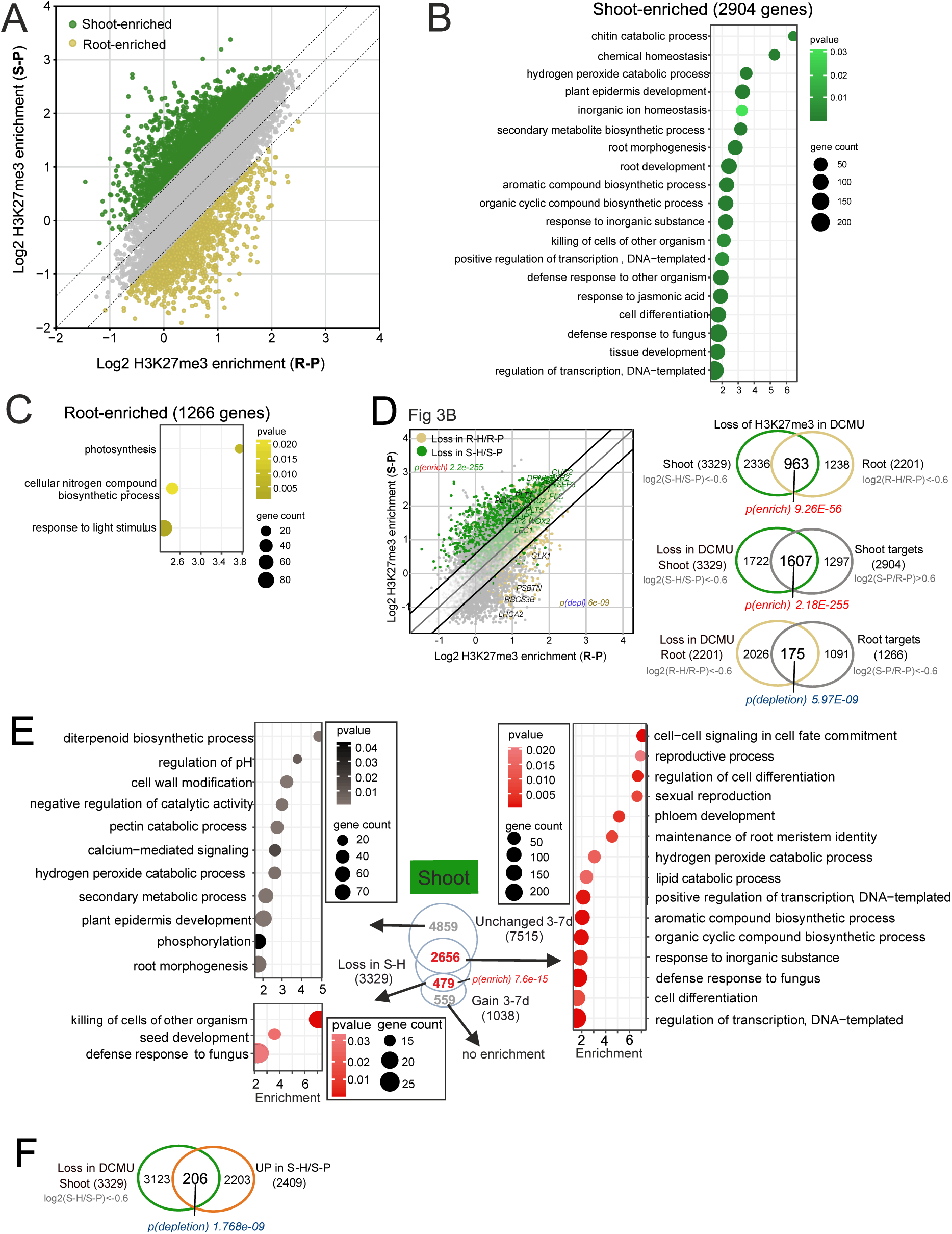
Shoot and Root specific target of H3K27me3 and impact of photosynthesis inhibition on shoot and root H3K27me3 target. (Related to Figure 3). **A)** H3K27me3/H3 enrichment of target genes in 7-day shoot (S-P) and root (R-P). Each dot represents a gene, all genes identified as H3K27me3-targets (T70) in either 7-dag shoot or 7-dag root are plotted. Green and ochre dots represent genes that are enriched for H3K27me3 in the shoot and root, respectively. **B)** GO analysis of genes enriched for H3K27me3 in the shoot only. **C)** GO analysis of genes enriched for H3K27me3 in the root only. BP categories are shown, GO display cutoff: fold enrichment > 2; p(Bonferroni) < 0.05. **D)** Extension to Figure 3B: H3K27me3/H3 enrichment of target genes in 7-dag shoot (S-P) and root (R-P). Each dot represents a gene; green and ochre dots represent genes that lose H3K27me3 in 7-dag heterotrophic shoot (S-H/S-P) and root (R-H/R-P), resp. p-values: significance of overlap (enrichment or depletion) between shoot (green) or root (ochre) genes and genes losing H3K27me3 in respective heterotrophic samples: Venn diagrams on the right. **E)** GO enrichment of H3K27me3 targets with unchanged or gain of H3K27me3 between 3- and 7-dag shoot (S-P3/S-P) that lose (red) or do not lose (grey) H3K27me3 in heterotrophic shoot (S-H). BP categories are shown, GO display cutoff: fold enrichment > 1.5; p(Bonferroni) < 0.05. **F)** Overlap between genes that lost H3K27me3 and that were upregulated in heterotrophic shoot compared to photoautotrophic shoot (S-H/S-P).

**Figure S5.**
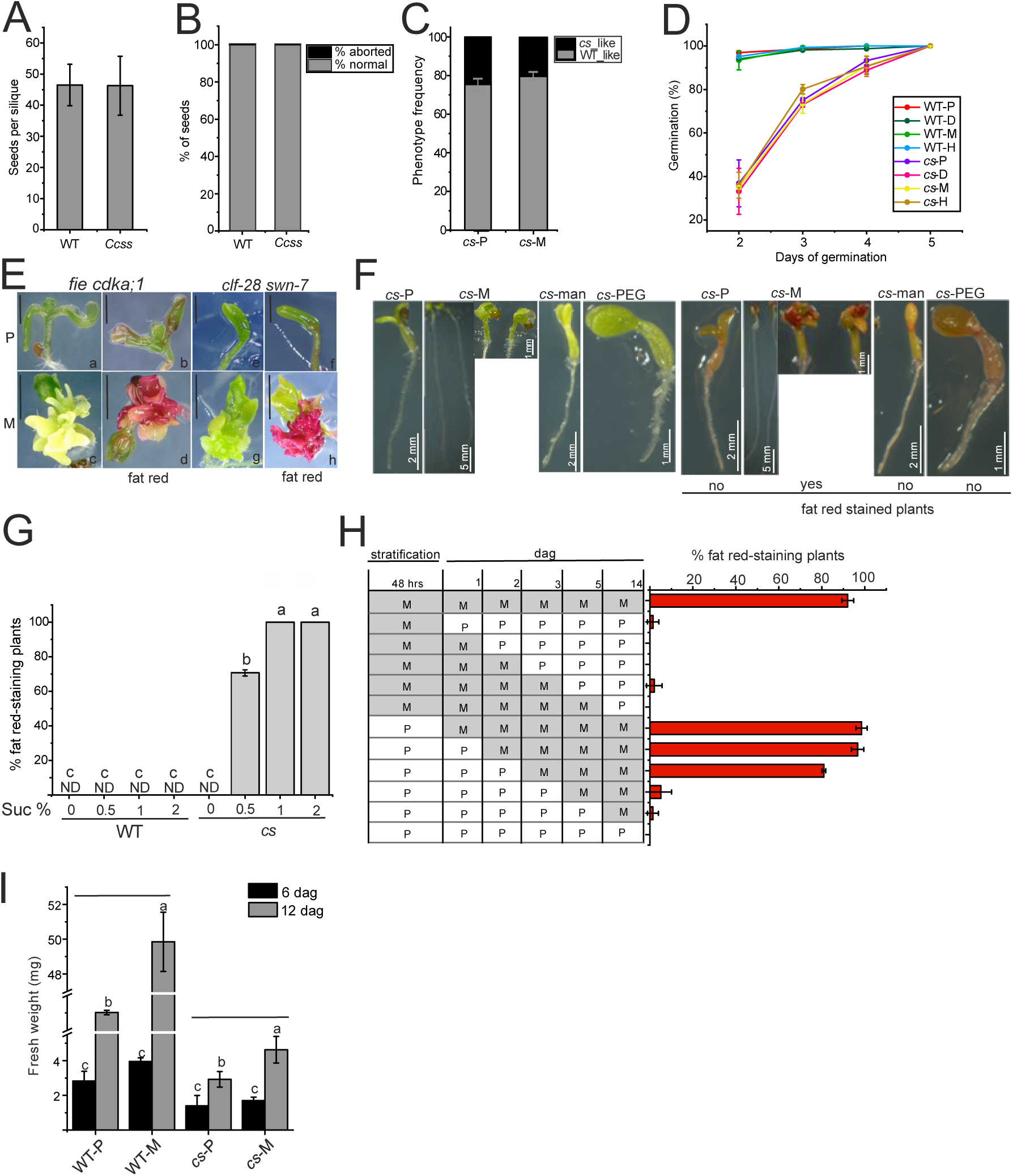
Absence of CLF and SWN does not reduce embryo or seed viability but is associated with delayed seed germination and sucrose-dependent TAG accumulation. (Related to Figure 4). **A)** Seed set in WT and *CLF/clf swn/swn* (*Ccss*). N = 58 WT and 80 *Ccss* siliques. **B)** Seed abortion rate in WT and *Ccss*. N = 4331 seeds (96 siliques, 17 individual plants) in WT and 6346 seeds (145 siliques, 15 individual plants) in *Ccss*. 0.23% and 0.34 % seeds aborted in WT and *Ccss*, resp. **C)** Penetrance of *clf swn* (*cs*) phenotype in 10-dag progeny of *Ccss* plants cultivated in mixotrophic (*cs*-M) or photoautotrophic (*cs*-P) conditions. The frequency does not deviate from the expected 25% (p 0.912, 0.991 for M, P respectively; χ2 test). Bars: mean ±SD; N = 6 biological replicates (130 - 260 seeds/replicate/growth condition). **D)** Cumulative percentage of germination of WT and *cs* in photoautrophic (WT-P, *cs*-P), mixotrophic (WT-M, *cs*-M), heterotrophic (WT-H, *cs*-H: +DCMU+suc) and +DCMU (WT-D, *cs*-D) conditions. N = 2 biological replicates (seeds/replicate/time point: 133 – 208 in WT and 31 - 62 in *cs*). **E)** Phenotype of *fie cdka;1* (a - d) and *clf-28 swn-7* (e - h) seedlings grown in photoautrophic (P) or mixotrophic (M) conditions. Fat red – plants stained by SudanRed7B to detect TAGs. Scale bar = 1 mm. N = 3 biological replicates (40- 50 *clf swn*). Penetrance of *fie* was approx. 2% in mixotrophic and not distinguishable in photoautotrophic conditions. **F)** Accumulation of TAGs in *clf swn* induced by sucrose but not by mock treatment, or osmotic controls including mannitol of PEG in the growth medium. 30 mM sugar was used. **G)** Penetrance of TAG-accumulating phenotype in populations of WT and *cs* plants grown in photoautotrophic (sucrose 0 %) and mixotrophic (sucrose 0.5%, 1% and 2%) growth conditions. Bars: mean ± SD; N = 3 biological replicates (25 - 59 *cs* or 200 WT 21-dag seedlings/replicate). Letters above bars: statistical significance at p < 0.05; two-way ANOVA with Bonferroni post hoc test. ND - not detected. **H)** Sucrose-induced TAG accumulation in *cs* is conditioned by the presence of sucrose within the first 2-3 days after seed germination induction. Timescale in the table indicates timepoints at which germinating seeds were transferred between cultivation plates with (M - mixotrophic) or without (P - photoautotrophic) 1% sucrose. Bars: mean ± SD; N = 3 biological replicates (20-30 14-dag *cs* plants/replicate). **I)** Biomass of WT and *cs* seedlings cultivated in photoautrophic (WT-P, *cs*-P) and mixotrophic (WT-M, *cs*-M) growth conditions for 6 and 12 days. Bars: mean ± SD; N = 4 biological replicates (5 seedlings/replicate were pooled). Letters above bars: statistical significance at p < 0.05; two-way ANOVA with Bonferroni post hoc test.

**Figure S6.**
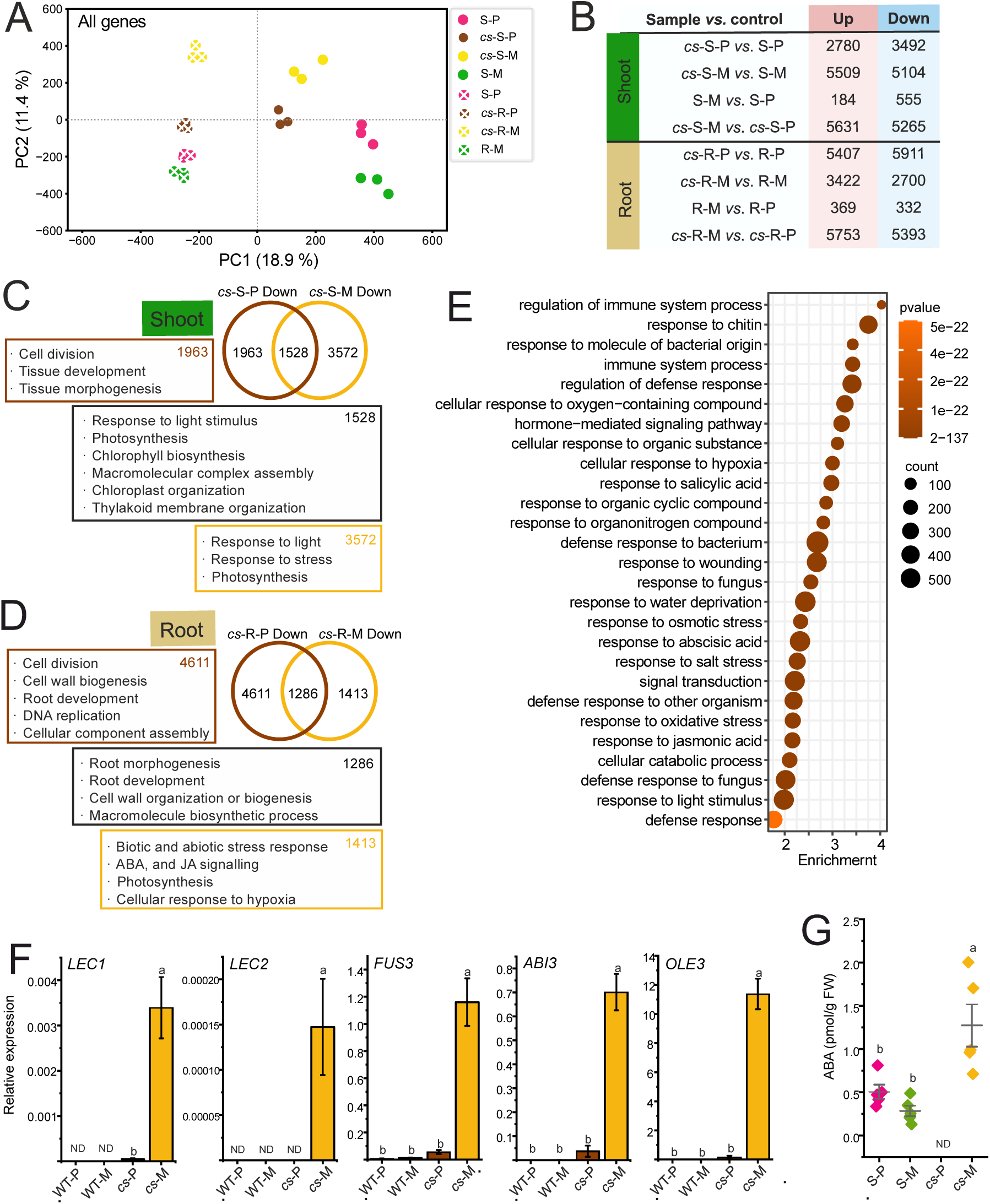
Transcriptomic analysis of the *clf swn* cultivated in photoautotrophic or mixotrophic conditions and ABA concentration in the samples. (Related to Figure 4). **A)** RNA-seq principal component analysis (PCA): RPKM values of all ARAPORT11 genes. Samples: wild-type (WT) and *clf swn* (*cs*); photoautotrophic (P) and mixotrophic (M); shoot (S) and root (R) tissues. **B)** Numbers of differentially expressed genes (DEGs) in the analysed sample comparisons. Cutoff: absolute log2FC > 0.6, FDR < 0.05 (commonly identified in EdgeR and DESeq2). **C)** Schematic representation of genes and enriched GO biological processes downregulated in *cs* photoautotrophic (*cs*-S-P) and mixotrophic (*cs*-S-M) shoot compared to respective WT shoot (S-P and S-M) samples. GO summary threshold: p(Bonferroni) < 0.05. **D)** Schematic representation of genes and enriched GO biological processes downregulated in *cs* photoautotrophic (*cs*-R-P) and mixotrophic (*cs*-R-M) root compared to respective WT root (R-P and R-M) samples. GO summary threshold: p(Bonferroni) < 0.05. **E)** Biological processes enriched among genes upregulated in photoautotrophic (*cs*-S-P) compared to mixotrophic (*cs*-S-M) *cs* shoot. GO display cutoff: p(Bonferroni) < 1e-21. **F)** Transcript abundances of *LEC1, LEC2, FUS3, ABI3* and *OLE3* determined by RT-qPCR wild type (WT) and *clf swn* (*cs*) under photoautotrophic (P) and mixotrophic (M) growth conditions. Bars: mean ± SD; N = 3 biological replicates. Letters above bars: statistical significance at p < 0.05; two-way ANOVA with Bonferroni post hoc test. **G)** Concentration of ABA in WT and *cs* shoot at 10 days. Bars: mean ± SD of 5 biological replicates. Letters above bars: statistical significance at p < 0.05; two-way ANOVA with Bonferroni post hoc test. ND - not detected.

**Figure S7.**
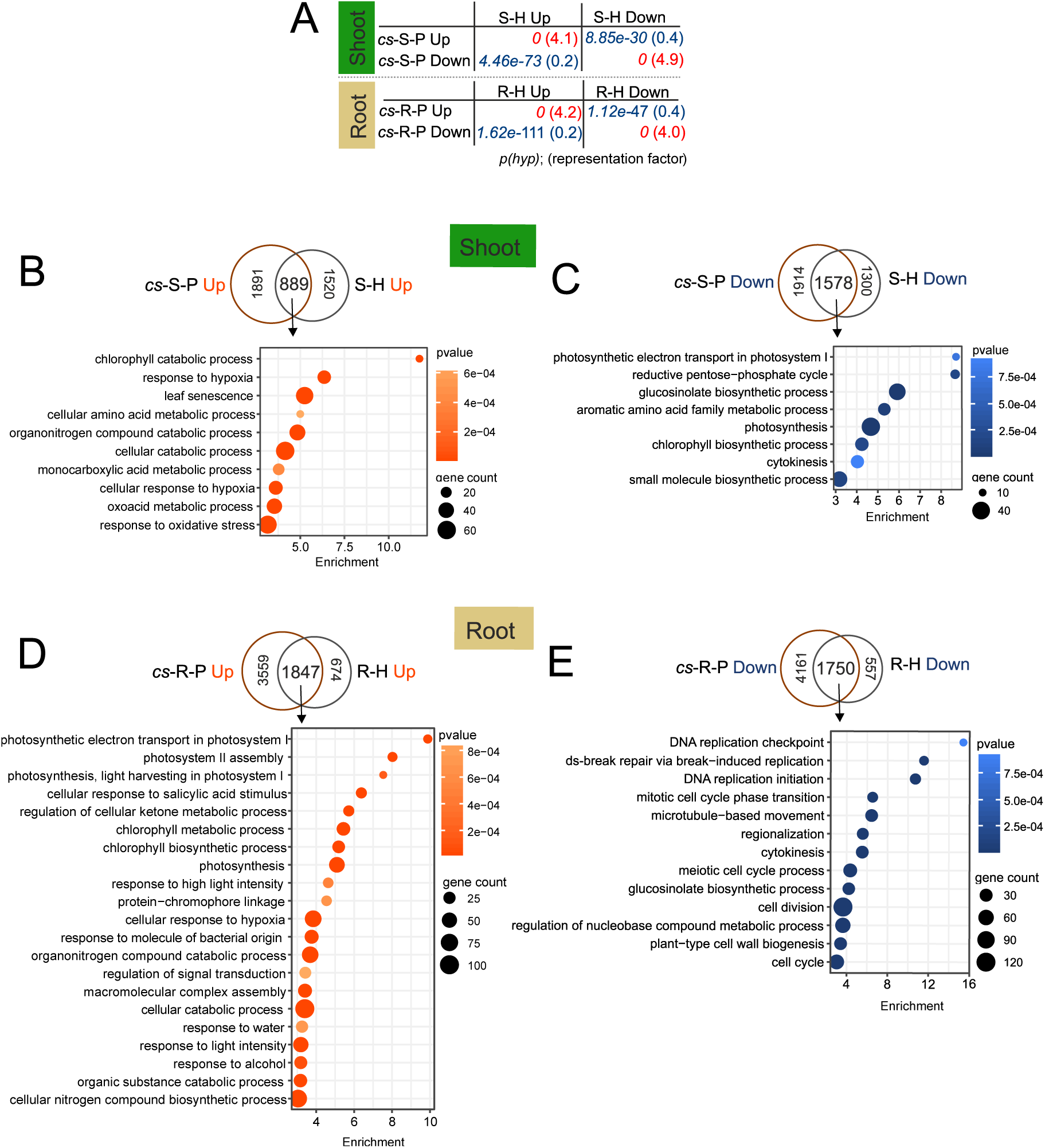
GO enrichment of biological processes commonly up- or down-regulated in photoautotrophic *cs* and DCMU-grown WT. (Related to Figure 4H). **A)** P-values of hypergeometric tests for overlaps and (representation factor) of dysregulated gene sets in photoautotrophic *cs* (*cs*-S/R-P) and heterotrophic WT (S/R-H) shoot (S) and root (R) tissues. Representation factor indicates enrichment (red font) or impoverishment (blue font) compared to expected. **B-E)** Biological processes enriched among: **B)** genes commonly upregulated in photoautotrophic *cs* shoot (cs-S-P) and heterotrophic, DCMU-grown WT shoots (S-H); **C)** genes commonly downregulated in photoautotrophic *cs* shoot (cs-S-P) and heterotrophic, DCMU-grown WT shoots (S-H); **D)** genes commonly upregulated in photoautotrophic *cs* root (cs-R-P) and heterotrophic, DCMU-grown WT roots (R-H); **E)** genes commonly downregulated in photoautotrophic *cs* root (cs-R-P) and heterotrophic, DCMU-grown WT roots (R-H). GO display cutoff: fold enrichment > 3; p(Bonferroni) < 0.001.

**Figure S8.**
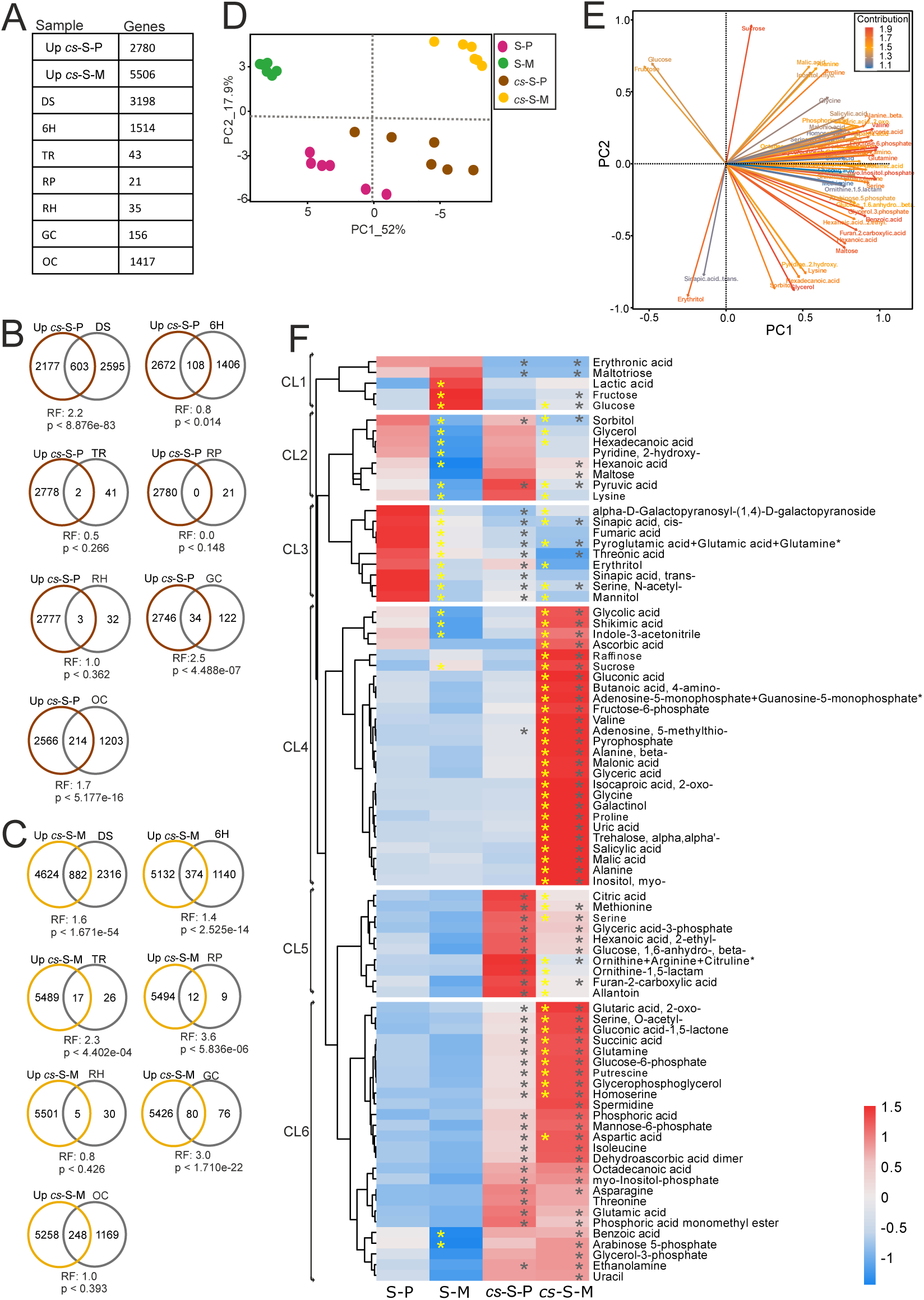
Comparison of transcriptomic and primary metabolome data. (Related to Figure 5). **A)** Summary of compared genes. The numbers of genes reported by Silva et al. 2016 correspond to clusters of genes that have maximum expression at a given developmental stage. **B)** and **C)** Overlap analysis of numbers of differentially expressed genes in **B)** photoautotrophic (*cs*-S-P) and **C)** mixotrophic (*cs*-S-M) *cs* shoot and genes within clusters described by Silva et al. 2016. P-values of hypergeometric tests for gene set overlaps are shown; RF: representation factor indicates enrichment (RF > 1) or impoverishment (RF < 1) compared to random overlap. DS - dry seed, 6H - six hours of imbibition, TR - testa rupture, RP - radicle protrusion, RH - root hair emergence stage, GC - cotyledon greening stage, OC - open cotyledons stage. **D)** Principal Component Analysis (PCA) of metabolite profiles. Relative abundance data were normalized to sample fresh weight and internal standard, log_10_-transformed, mean-centred, and auto-scaled prior to PCA generation. Biplot of PC1 and PC2 with respective % of represented variance. **E)** Contribution plot of 55 metabolites contributing to PC1 and PC2. **F)** Heatmap of relative abundances of 85 metabolites in WT photoautotrophic (S-P) and mixotrophic (S-M) shoot and *cs* photoautotrophic (*cs*-S-P) and mixotrophic (*cs*-S-M) shoot samples (Means, n = 5 biological replicates, red-blue colour coded z-score normalized by row). Metabolites are sorted into 6 clusters by hierarchical cluster analysis to the left. Asterisks (*): statistical significance at p < 0.05 (adjusted) based on ANOVA and Tukey’s HSD test. Black asterisks indicate significant difference among genotypes (*cs*-S-P vs. S-P and *cs*-S-M vs. S-M), yellow asterisks indicate significant differences induced by mixotrophy (S-M vs. S-P and *cs*-S-M vs. *cs*-S-P). Clusters (CL) 1-6 correspond to Figure. 5C.

**Figure S9.**
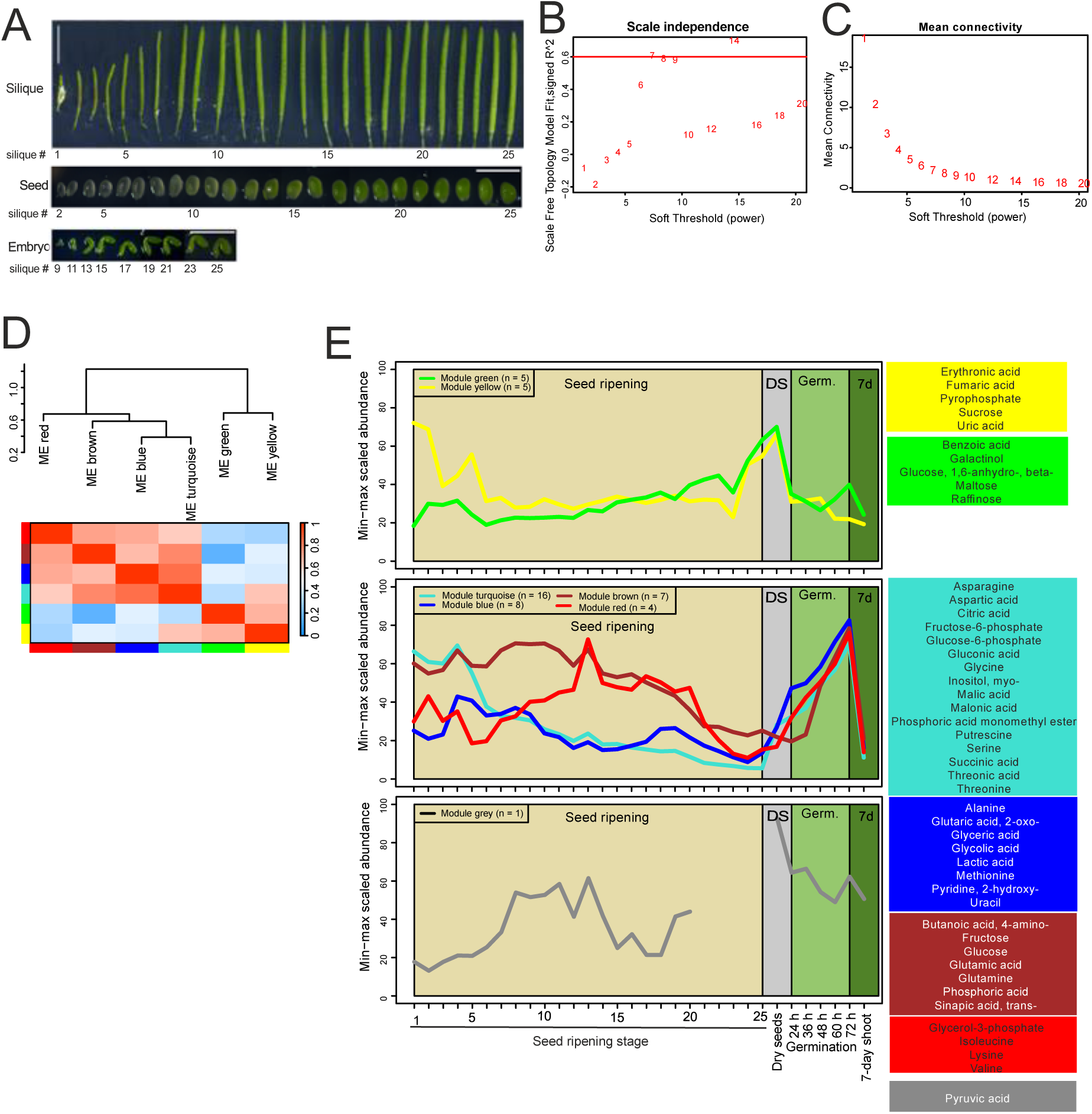
Weighted correlation network analysis of primary metabolome profiles from 25 seed maturation stages, dry seeds, and 5 seed germination stages (Related to Figure 5). **A)** 25 seed maturation stages of WT used for the profiling of the primary metabolome. Whole siliques were harvested at stages 1 – 6; seeds were harvested at stages 7 – 25. Seed maturation stages correspond to embryo development stages in the harvested material as shown by the bottom panel. **B-E)** Weighted correlation network analysis of 53 metabolites identified in samples of WT seed maturation stages. These metabolites were robustly present in all samples analyzed by the current study. **B)** and **C)** Optimization of the soft-thresholding power β parameter for the weighted metabolites correlation network analysis(Langfelder and Horvath, 2008). Weighted networks rely on metabolite-adjacencies, which correspond to the absolute correlations exponents of the soft-thresholding power β. The value β = 7 was chosen to obtain a network with Scale-Free Topology properties (Scale Free Topology Model Fit R2 ≥ 0.6) **(B)** and high connectivity **(C)**. **D)** Adjacency values were used to compute the Topological Overlap Matrix (TOM). Metabolites were clustered into modules (ME) based on the (1-TOM) distance matrix, by hierarchical clustering with average linkage distance comparison between clusters. The number of modules was determined by Dynamic Hybrid tree cut (dendrogram cut height for module merging = 0.25). **E)** Metamodules were identified based on the results of hierarchical clustering of the eigenmetabolites representing each ME, considering the (1-TOM) distance.

**Figure S10.**
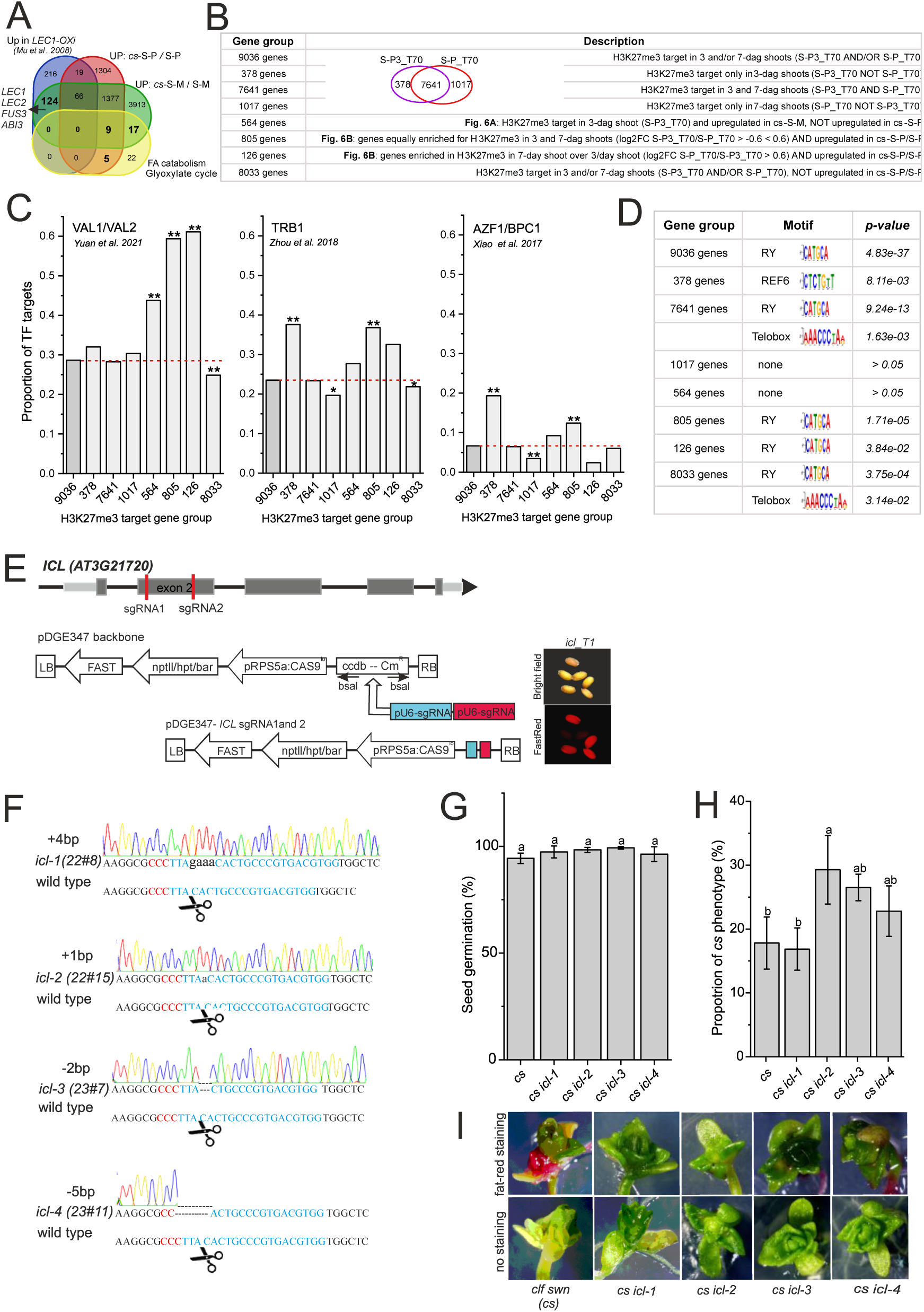
PRC2 recruitment modules involved in seed-to-seedling transition and CRISPR/Cas9-mutagenesis of *ICL* in *cs*. (Related to Figure 6). **A)** Overlap of genes upregulated in photoautotrophic (*cs*-S-P) or mixotrophic (*cs*-S-M) *cs* shoot compared to respective WT shoot controls, genes upregulated in *LEC1*- overexpressing plants (*LEC1-Oxi* - *pER8-LEC1*) compared to wild type (Mu et al., 2008) and genes involved in metabolic processes marking seed germination (the list of genes was downloaded from plant metabolic network (PMN: https://plantcyc.org/: fatty acid β-oxidation (BioCyc ID: PWY-5136), glyoxylate cycle (BioCyc ID: GLYOXYLATE-BYPASS), TCA cycle (BioCyc ID: PWY-5690), gluconeogenesis (BioCyc ID: GLUCONEO-PWY)). *LEC1-Oxi-*upregulated genes showed a significant overlap (hypergeometric test) with *cs*-M (p < 6.27e-43) and *cs*- P (p < 3.95e-14). *cs*-M (p < 2.62e-05) but also *cs*-P (p < 0.00146) showed an overlap with genes involved in seed germination-related metabolic pathways while none of these genes were upregulated in *LEC1-Oxi*. **B)** Groups of H3K27me3 target genes analysed in C and D. **C)** Proportion of transcription factor target genes among H3K27me3 targets. All H3K27me3 targets identified in 3-dag and/or 7-dag shoot (9036 genes) were used as background (dark grey column, indicated by red dashed line). Significant enrichment or depletion to background: *p < 0.01, **p < 0.0001; two-sided Fisheŕs exact test. **D)** Polycomb response element (PRE)/REF6-motif enrichment among H3K27me3 targets. **E - I)** Generation and analysis of CRISPR/Cas9 *cs icl* lines. **E)** *ICL* (*AT3G21720*) locus and the positions targeted by the two guide RNAs (sgRNAs). *CRISPR/Cas9* construct used (Stuttmann et al., 2021) to transform *CLF/clf swn/swn* (*Ccss)* plants. **F)** Sanger sequencing chromatogram and sequence of *ICL* in the four independent *clf swn* (*cs*) *icl* lines used. The gRNA sequence is shown in blue colour, the PAM sequence is shown as 3 bps in red colour, and scissors indicate the Cas9 cut site. Numbers in bp with (-) or (+) indicate deletion or insertion, respectively. **G)** Seed germination in the progeny of *Ccss* plants is not affected by mutation in *ICL*. **H)** Frequency of *cs* phenotype is not reduced by *ICL* mutation in the progeny of *Ccss icl* plants. **I)** Representative images of *cs icl* plants analysed by fat-red staining – images complement statistics in Figure 6F. T3 Cas9-negative plants in mixotrophic (1% sucrose) conditions were analysed in G)-I). Bars: mean ± SD; N = 3 biological replicates (95-105 seeds/replicate). Letters above bars: statistical significance at p < 0.05; one-way ANOVA with Bonferroni post hoc test.

